# NUCLEAR FACTOR Y, subunit C (NF-YC) transcription factors are positive regulators of photomorphogenesis in *Arabidopsis thaliana*

**DOI:** 10.1101/067462

**Authors:** Zachary A. Myers, Roderick W. Kumimoto, Chamindika L. Siriwardana, Krystal K. Gayler, Jan R. Risinger, Daniela Pezzetta, Ben F. Holt

## Abstract

Recent reports suggested that NF-Y transcription factors are positive regulators of skotomorphogenesis in *Arabidopsis thaliana*. Three *NF-YC* genes (*NF-YC3, NF-YC4*, and *NF-YC9*) are known to have overlapping functions in photoperiod dependent flowering and previous studies demonstrated that they interact with basic leucine zipper (bZIP) transcription factors. This included ELONGATED HYPOCOTYL 5 (HY5), which has well-demonstrated roles in photomorphogenesis. Similar to *hy5* mutants, we report that *nf-yc3 nf-yc4 nf-yc9* triple mutants failed to inhibit hypocotyl elongation in all tested light wavelengths. Surprisingly, *nf-yc3 nf-yc4 nf-yc9 hy5* mutants had synergistic defects in light perception, suggesting that NF-Ys represent a parallel light signaling pathway. As with other photomorphogenic transcription factors, *nf-yc3 nf-yc4 nf-yc9* triple mutants also partially suppress the short hypocotyl and dwarf rosette phenotypes of CONSTITUTIVE PHOTOMORPHOGENIC 1 (*cop1*) mutants. Thus, our data strongly suggest that NF-Y transcription factors have important roles as positive regulators of photomorphogenesis, and in conjunction with other recent reports implies that the NF-Y are multifaceted regulators of early seedling development.

**Author Summary:** Light perception is critically important for the fitness of plants in both natural and agricultural settings. Plants not only use light for photosynthesis, but also as a cue for proper development. As a seedling emerges from soil it must determine the light environment and adopt an appropriate growth habit. When blue and red wavelengths are the dominant sources of light, plants will undergo photomorphogenesis. Photomorphogenesis describes a number of developmental responses initiated by light in a seedling, and includes shortened stems and establishing the ability to photosynthesize. The genes regulating photomorphogenesis have been studied extensively, but a complete picture remains elusive. Here we describe the finding that NUCLEAR FACTOR-Y (NF-Y) genes are positive regulators of photomorphogenesis - i.e., in plants where NF-Y genes are mutated, they display some characteristics of dark grown plants, even though they are in the light. Our data suggests that the roles of NF-Y genes in light perception do not fit in easily with those of other described pathways. Thus, studying these genes promises to help develop a more complete picture of how light drives plant development.

## Introduction

Plants utilize multiple properties of light, such as intensity, quality, and direction, to guide growth and development (1). The effects of light on plant development are exemplified by the transition of seedlings from dark growth (where they exhibit skotomorphogenesis) to light growth (photomorphogenesis). This transition is crucial for plant viability and is characterized by the inhibition of hypocotyl elongation, the expansion of cotyledons, and the accumulation of photosynthetic pigments. In *Arabidopsis thaliana*, several different classes of photoreceptors mediate light perception, including the phytochromes which perceive red and far red light, cryptochromes, phototropins, and LOV (Light, Oxygen, or Voltage) domain proteins, which are blue light receptors, and UV RESISTANCE LOCUS 8, the most-studied morphogenic photoreceptor for UV-B light (2–6). Of these receptors, the photomorphogenic transition is primarily controlled through the actions of the phytochromes and the cryptochromes (7). Through their combined actions, signaling cascades are initiated that significantly modify the expression of at least two thousand genes in *Arabidopsis* (8).

While sustained photomorphogenic growth requires the actions of multiple phytochromes and cryptochromes, the initial signaling cascade is established primarily through phyA, which accumulates to high levels in darkness (9). Upon activation by far red light, phyA is imported into the nucleus through interactions with FAR RED ELONGATED HYPOCOTYL 1 (FHY1) and FHY1-LIKE (FHL) (10). The physical interaction of phyA with FHY1/FHL is also necessary for phyA to bind further downstream transcription factors that regulate light signaling (11, 12). The function of phyA, as well as the photomorphogenic downstream transcription factors, is modulated at multiple levels, including through phyA-mediated protein phosphorylation and targeted, proteasome-mediated degradation. The ubiquitination and targeting of many photomorphogenic proteins for proteasome degradation is regulated through the actions of CONSTITUTIVE PHOTOMORPHOGENESIS 1 (COP1). In the light and in response to the initiation of phytochrome-mediated signal transduction, COP1 protein is excluded from the nucleus, allowing the accumulation of photomorphogenesis-promoting transcription factors (13–15).

One of the most significant targets of COP1 is HY5, a relatively small bZIP transcription factor that regulates photomorphogenesis by activating a large number of further downstream transcription factors (16–18). HY5 has also been identified as an integrator of pathways not directly related to light signaling, including hormone signaling (abscisic acid (ABA) and brassinosteroids), apoptosis, and temperature acclimation (19–22). Unlike many other genes involved in the light signaling cascade, the phenotypes of *hy5* mutants are not wavelength-specific (18, 23). Other COP1 targets, including the bHLH transcription factor LONG HYPOCOTYL IN FAR-RED 1 (HFR1) and the MYB transcription factor LONG AFTER FAR-RED LIGHT 1 (LAF1), function in a wavelength-dependent manner; while both *hfr1* and *laf1* mutants have reduced responses to far red light, only *hfr1* has a visible phenotype in blue light, and neither mutant exhibits phenotypes in red or white light (24–28). Further elucidation of the light-signaling cascade has revealed a handful of other transcription factors whose function is necessary for normal photomorphogenic growth and are also regulated by COP1, including the B-box (BBX) containing proteins SALT TOLERANCE HOMOLOG 2 (STH2/BBX21) and LIGHT REGULATED ZINC FINGER1/STH3/BBX22, the bHLH proteins PHYTOCHROME RAPIDLY REGULATED 1 (PAR1) and PAR2, and the *Mutator* transposase-like FAR-RED ELONGATED HYPOCOTYL 3 (FHY3) and FAR-RED IMPAIRED RESPONSE 1 (FAR1) (29–32). Thus, light perception, and the associated photomorphogenic signaling cascades, converges at a small suite of transcription factors just downstream of a COP1-mediated hub. From these terminal transcription factors it appears that the signal cascade immediately fans out to thousands of light-regulated genes (33, 34).

While significant progress has been made identifying and characterizing transcription factors functioning at this COP1-mediated hub, there are likely to be undiscovered pieces in this puzzle. No combination of downstream transcription factor mutants that can phenocopy the *phyA* mutant has been identified – e.g., when grown in far red light, the hypocotyls of *hy5 hfr1 laf1* triple mutants are ~60% as long as the *phyA* mutant (35). While this triple mutant has significantly longer hypocotyls than any of the single mutants or double mutant combinations, residual far red light perception is clearly still present. This is in contrast to *fhy1 fhl* double mutants which appear phenotypically identical to both *phyA* mutants and dark grown plants for hypocotyl elongation (10, 36, 37). One explanation is that the downstream transcription factor components are already known, but the right combination of mutations has yet to be assembled in a single genotype. For example, as with *hy5 hfr1 laf1* plants, *hy5 sth2 sth3* mutants are also additively defective in light perception (30), but no *hy5 hfr1 laf1 sth2 sth3* higher order mutant has been reported. Alternatively, additional transcription factor components may remain unknown. Following two recent publications ((38, 39), we report here additional strong evidence for the involvement of NUCLEAR FACTOR Y (NF-Y) transcription factors in light perception.

NF-Y transcription factors consist of three unique proteins, called NF-YA, NF-YB, and NF-YC, and each is encoded by a small family of ~10 genes in Arabidopsis (this expansion is mirrored in other sequenced plant species, including monocots and dicots; (40, 41)). None of the NF-Y subunits is thought to regulate transcription independently; instead, the mature NF-Y transcription factor is composed of one of each subunit type and all three subunits contribute to DNA binding. NF-YB and NF-YC initially form a dimer in the cytoplasm that translocates to the nucleus where a trimer is formed with NF-YA (42–46). Thus, regulation of any one NF-Y subunit can alter the function of the entire complex. Following nuclear assembly of the mature complex, NF-Ys bind DNA at CCAAT-containing *cis* regulatory elements and are typically positive regulators of gene expression (47). Although the generalized characterization of NF-Ys (largely from animal and yeast systems) describes them as binding DNA in the proximal regions of promoters, recent data suggests that they also bind more distal regions of promoters to regulate gene expression (48, 49). In the animal lineage, each NF-Y is usually encoded by only one or two genes and the functional consequences of expanded *NF-Y* gene families in the plant lineage remains only modestly explored. Nevertheless, much progress has been made in recent years describing the roles of individual NF-Y subunits in the control of specific processes, especially the control of photoperiod-dependent flowering through interactions with CONSTANS (CO) (50–52), various functions in the development of nitrogen-fixing root nodules in legume species (53–55), and abscisic acid signaling during germination and early seedling establishment, often mediated by interactions with bZIP transcription factors (38, 39, 56–58).

Relevant to NF-Y roles in photomorphogenesis and light perception, little is currently known. However, NF-YA5 and NF-YB9 were previously implicated in regulating blue light-dependent transcript accumulation for LIGHT-HARVESTING CHLOROPHYL A/B BINDING PROTEIN (59) and NF-Y complexes were also shown to bind and regulate the expression of the spinach photosynthetic gene AtpC (60). Further, the promoters of a number of light signaling components were bound by LEC1/NF-YB9 (LEAFY COTYLEDON 1 (61, 62)) in chromatin immunoprecipitation experiments, including light harvesting and chlorophyll binding proteins (e.g., LHCA1 and LHCB5) and transcriptional regulators of light perception (e.g., HY5, HY5 HOMOLOG (HYH), and HFR1) (38). Finally, alterations in hypocotyl elongation resulting from both *NF-YB* loss of function and inducible overexpression have been observed (38, 63), including the recent report that LEC1/NF-YB9 regulates skotomorphogenesis through physical interaction with PHYTOCHROME-INTERACTING FACTOR 4 (39).

Previous work in our lab identified physical interactions between NF-YC and HY5, as well as other bZIP proteins (57). Here we extend these initial observations to show that these same NF-YC proteins (NF-YC3, 4, and 9) are broad spectrum regulators of light perception. Interestingly, in the same way that HY5, HFR1, and LAF1 can physically interact, but still appear to signal through independent pathways, *hy5 nf-yc* mutants also show additive - even synergistic - light perception defects. This manuscript characterizes several photomorphogenesis-related phenotypes of NF-YC mutants and proposes that the NF-Y complexes constitute a novel component of the light signaling cascade, functioning at least partially independent of HY5, HFR1, and LAF1. We further demonstrate that *nf-yc* mutants can partially suppress several *cop1* mutant phenotypes and that proteasome regulation of NF-Y complexes during light perception is mitigated through NF-YA subunits. Similar to the multiple regulatory roles of HY5 in light perception and abscisic acid (ABA) signaling, our cumulative research on these three NF-Y proteins demonstrates that they have essential roles in photoperiod-dependent flowering, ABA perception, and light perception.

## Results

### Inhibition of hypocotyl elongation in short day, white light, and several individual light wavelengths requires NF-YC

We initially observed slightly elongated hypocotyls in plate grown *nf-yc3-1 nf-yc4-1 nf-yc9-1* triple mutants (hereafter *nf-yc triple*, (51)). These visual differences primarily appeared in plants grown for shorter day lengths. To quantify these observed differences, we compared *nf-yc triple* mutants to their parental Columbia (Col-0) ecotype under continuous white light (cWL), short day (SD, 8hrs light/16hrs dark), and continuous dark (cD) conditions (Fig 1A–D). While continuous light and continuous dark grown *nf-yc triple* mutants were not significantly different from Col-0, short day grown seedlings had moderately elongated hypocotyls (~50–60% longer, Fig 1B–C). To de-convolute the contributions of individual *NF-YC* genes, we additionally examined hypocotyl elongation for the six possible single and double mutants from the three mutant alleles. Modest differences were observed for only the *nf-yc3 nf-yc9* double mutant (Fig 1B), although we note that this mutant phenotype was inconsistent in additional experiments. Overall, the data suggested that *NF-YC3, NF-YC4, and NF-YC9* were collectively necessary for the suppression of hypocotyl elongation. Supporting the genetic data showing overlapping functions, all three genes were strongly expressed in the hypocotyl with peak expression in the vascular column (Fig S1).

**Figure 1.**
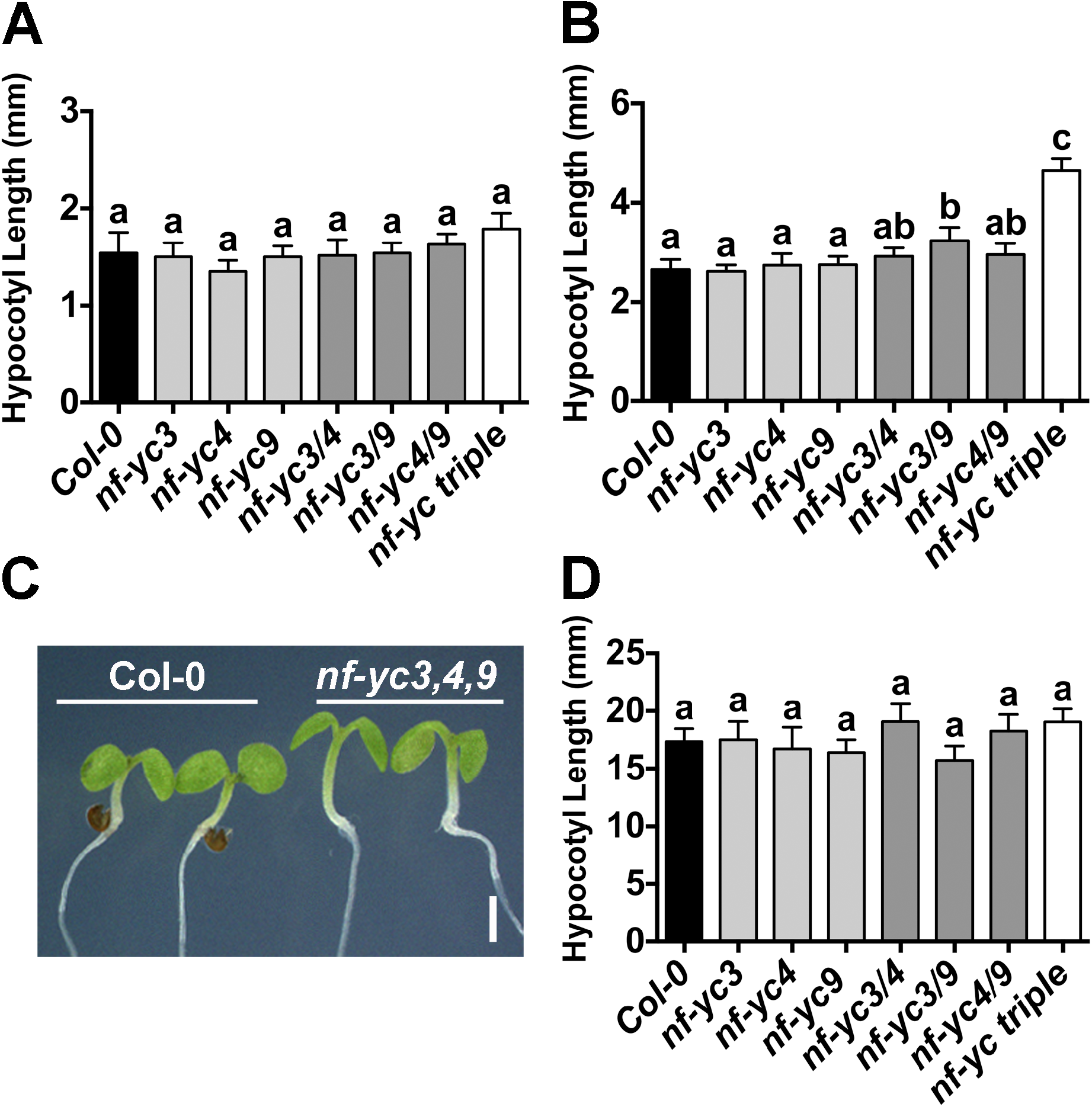
*NF-YC3, 4*, and *9* contribute redundantly to suppression of hypocotyl elongation in white light. Hypocotyl lengths are shown for plants grown for five days on B5 media in **A**) cWL, **B-C**) SD, or **D**) cD conditions. Statistically significant differences (or lack thereof) are represented by lettering above bars (error bars 95% confidence interval). Statistical differences were determined by ANOVA (P<0.01) and subsequent multiple comparisons by either Tukey’s (cWL) or Dunnett’s (SD, cD) procedures. Scale bar in C) represents 2mm.

To determine whether the hypocotyl elongation defects were wavelength specific, we additionally examined the same suite of mutants grown in continuous blue (cB), far red (cFR), and red (cR) light conditions (Fig 2A–C). In cFR conditions, no significant differences were observed. However, in cR and cB light the *nf-yc triple* mutants were ~50% longer than Col-0. Additionally, significant hypocotyl elongation defects were observed in some single and double mutants (ranging from ~18–29% longer that Col-0). As with the short day white light measurements, differences in the single and double mutants in cB and cR light were less robust between repeated experiments than for the *nf-yc triple* mutants. Interestingly, longer hypocotyls were always associated with the presence of the *nf-yc9* mutant allele – somewhat unexpected as the *nf-yc3-1* and *nf-yc4-1* alleles are strong knockdowns while the *nf-yc9-1* allele retains ~20–25% normal expression levels (Fig S2 and (51)). Collectively, these data demonstrate that NF-YCs are broad spectrum regulators of light perception.

**Figure 2.**
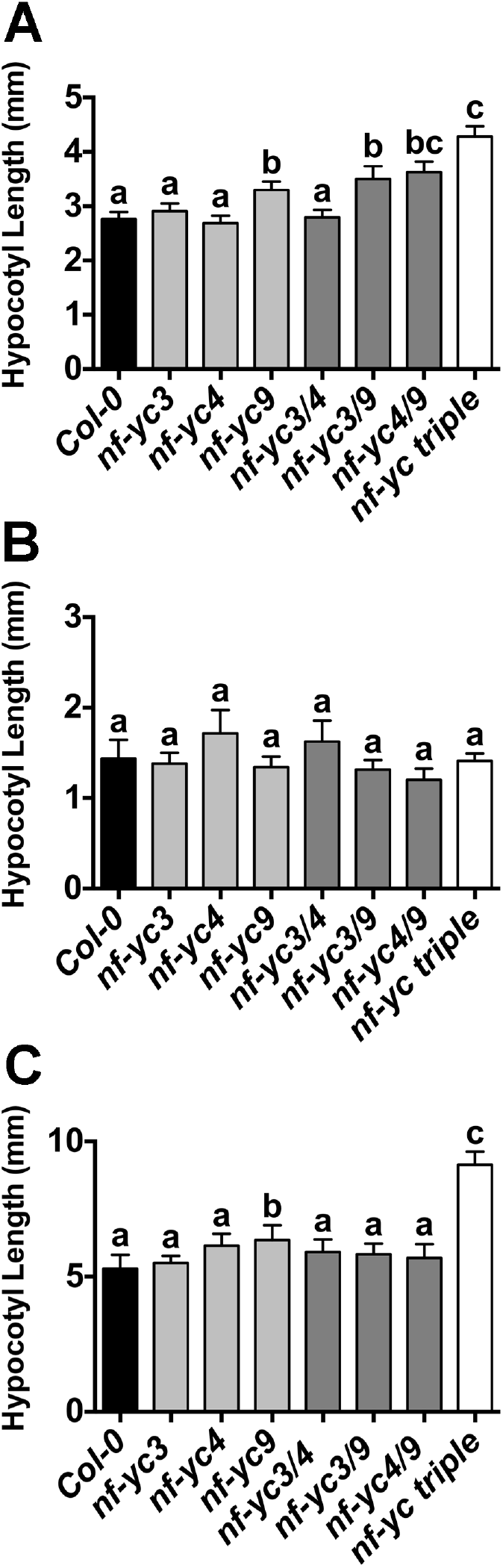
NF-YC3, 4, and 9 are necessary for suppression of hypocotyl elongation in both cB and cR light. Hypocotyl lengths are shown for five day old plants grown on B5 media in **A**) cB (38μmol m^−2^ s^−1^), **B**) cFR (5μmol m^−2^ s^−1^), and **C**) cR (6μmol m^−2^ s^−1^). Statistically significant differences between groups (or lack thereof) are represented by lettering above bars (error bars 95% confidence interval). Statistical differences were determined by standard ANOVA (p<0.01) when variances were not significantly different (cFR and cR) and Kruskal-Wallis ANOVA (non-parametric test, p<0.05) when variances were unequal (cB). Subsequent multiple comparisons were performed by either Tukey’s (cFR, cR) or Dunn’s (cB) procedures, respectively.

### FLIM-FRET Analysis Confirms the HY5 by NF-YC9 Physical Interaction

NF-Y complexes are known to associate with bZIP transcription factors in both plants and animals (46, 64–66). Relevant to light perception, we previously reported a modest yeast two-hybrid (Y2H) interaction between NF-YC4 and NF-YC9 with HY5 and a noninteraction with NF-YC3, although we assumed the inability to detect an NF-YC3 interaction was likely due to its autoactivation problems in the Y2H system (57). To further confirm this Y2H data, we performed transient interaction assays in *Nicotiana benthamiana* (Fig 3A–C). We utilized fluorescence lifetime imaging (FLIM) and fluorescence recovery after photobleaching (FRAP) to detect fluorescence resonance energy transfer (FRET) between epitope-tagged NF-YC and HY5 proteins. In these experiments, HY5 or NF-YB2 (positive control for interaction with NF-YC) were translationally fused to enhanced yellow fluorescent protein (YFP, (67)) and assayed for FRET against NF-YC3, 4, and 9 fused to modified cerulean 3 (mCer3, (68)). By comparing fluorescence lifetimes of the donor (mCer3), pre- and post-photobleaching of the acceptor (YFP), we could infer whether or not chosen protein pairs were closely physically associated. Direct physical interaction between proteins was indicated by a significant increase in the lifetime of mCer3 upon YFP photobleaching (69). Fluorescence recovery of both mCer3 and YFP was monitored during acceptor photobleaching, and was used as an internal control for balancing the destruction of YFP signal and the preservation of mCer3 signal during experimentation (Fig 3B). After identifying a proper photobleaching regimen, in pairs of known interacting proteins we observed that mCer3 signal would increase over the course of the initial photobleaching event, but not over subsequent treatments (Fig 3B). This was consistent with what is expected when observing FRET, as a significant majority of the acceptor (YFP) is destroyed in the initial photobleaching event, and further photobleaching events have a reduced effect on the already diminished pool of YFP.

**Figure 3.**
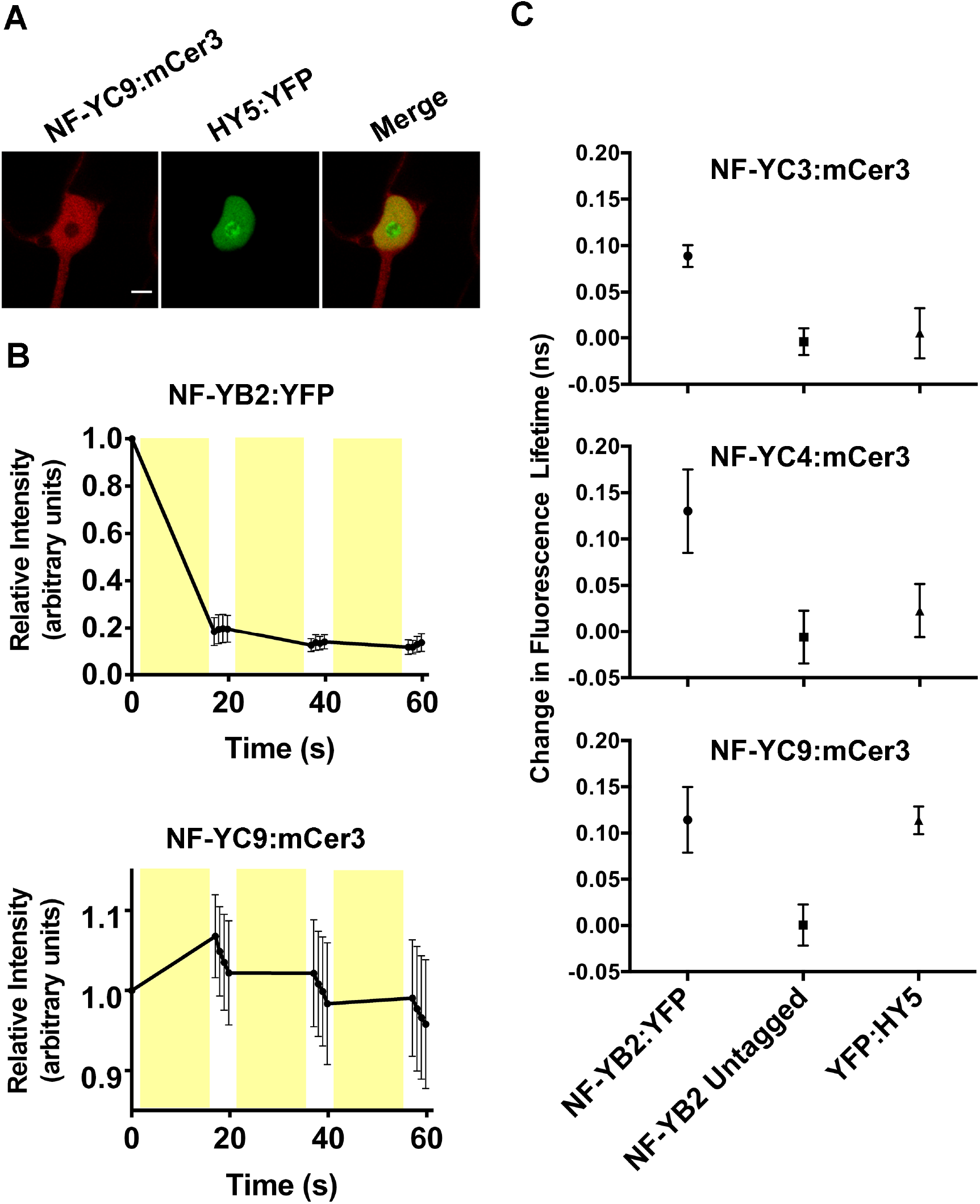
FRET-FLIM analysis shows a strong NF-YC9 by HY5 physical interaction. FRET experiments were conducted in tobacco leaves through transient 35S-driven overexpression of NF-YCs tagged with mCer3 and HY5 or NF-YB2 tagged with YFP. **A**) Nuclei expressing both mCer3 and YFP constructs were assayed for FRET through both FRAP and FLIM. **B**) A FRAP curve representative of a positive FRET result between two known interacting proteins, NF-YB2 and NF-YC9. Fluorescence intensity was calculated relative to the prephotobleached intensity of each fluorescent protein. Yellow bars represent the timing of photobleaching events. **C**) FLIM was employed to detect FRET through lifetime measurements before and after acceptor photobleaching (FRAP) within the same nucleus. Each point is an independent combination of mCer3- and YFP-tagged proteins, and represents the shift in fluorescent lifetime elicited by acceptor photobleaching. Scale bar in A) represents 5 μm. Error bars in B-C) represent 95% CI with an n 3.

As a positive interaction control, we initially tested NF-YB2:YFP by NF-YC3, 4, and 9:mCer3 and were able to consistently detect significantly increased mCer3 fluorescence lifetimes after YFP photobleaching (Fig 3C). This is consistent with previous publications showing strong Y2H and *in vivo* NF-YB by NF-YC interactions (51, 52, 70–72). As a negative control for each interaction test, we demonstrated that when NF-YB2 lacked the YFP fusion, mCer3 lifetimes were not altered after a photobleaching treatment (Fig 3C). Substituting HY5:YFP for NF-YB2:YFP demonstrated that NF-YC9 could consistently physically interact with HY5; however, no FLIM-FRET interaction was detected between NF-YC3 or NF-YC4 and HY5. Thus, it remains possible that HY5 only interacts with a subset of the light perception-regulating NF-YC proteins described here (see Discussion).

### NF-YC and HY5 genetically interact to suppress hypocotyl elongation in white light

With the knowledge that at least some NF-YCs can physically interact with HY5, we generated *nf-yc triple hy5* mutants and examined them for hypocotyl elongation phenotypes in both SD and cWL (Fig 4A–C). Surprisingly, in SD conditions the *nf-yc triple hy5* mutants were considerably longer than either parental mutant line, suggesting that the previously observed NF-YC roles in hypocotyl elongation were at least partially independent of HY5. Even more striking was the strongly synergistic increase in hypocotyl elongation in cWL in the *nf-yc triple hy5* mutants over both mutant parents (Fig 4C). Compared to parental Col-0, dark grown plants showed no differences in hypocotyl elongation for any of the mutant genotypes (Fig 4D). Rescue assays confirmed that each gene (*NF-YC3, 4, 9* and *HY5*) was capable of significantly suppressing the *nf-yc triple hy5* elongated hypocotyl phenotype (Fig S3). Collectively, these data demonstrated that the presence of *HY5* masked the effects of the *nf-yc triple* mutant on hypocotyl elongation, especially in cWL conditions. These results are not trivially explained by cross regulation between NF-YC and HY5 as their transcription levels are only altered in their own mutant backgrounds (Fig S2). Finally, because some commercial white light sources contain contaminating UV radiation, we additionally examined hypocotyl lengths of cWL-grown plants grown under Mylar to filter out UV light. In accordance with previous work, *hy5* mutants were longer in the absence of UV (73); however, no difference was detected in the *nf-yc triple* mutant, and while not statistically significantly different, the minor difference in *nf-yc triple hy5* mutants can be completely accounted for by the loss of *HY5* (Fig S4).

**Figure 4.**
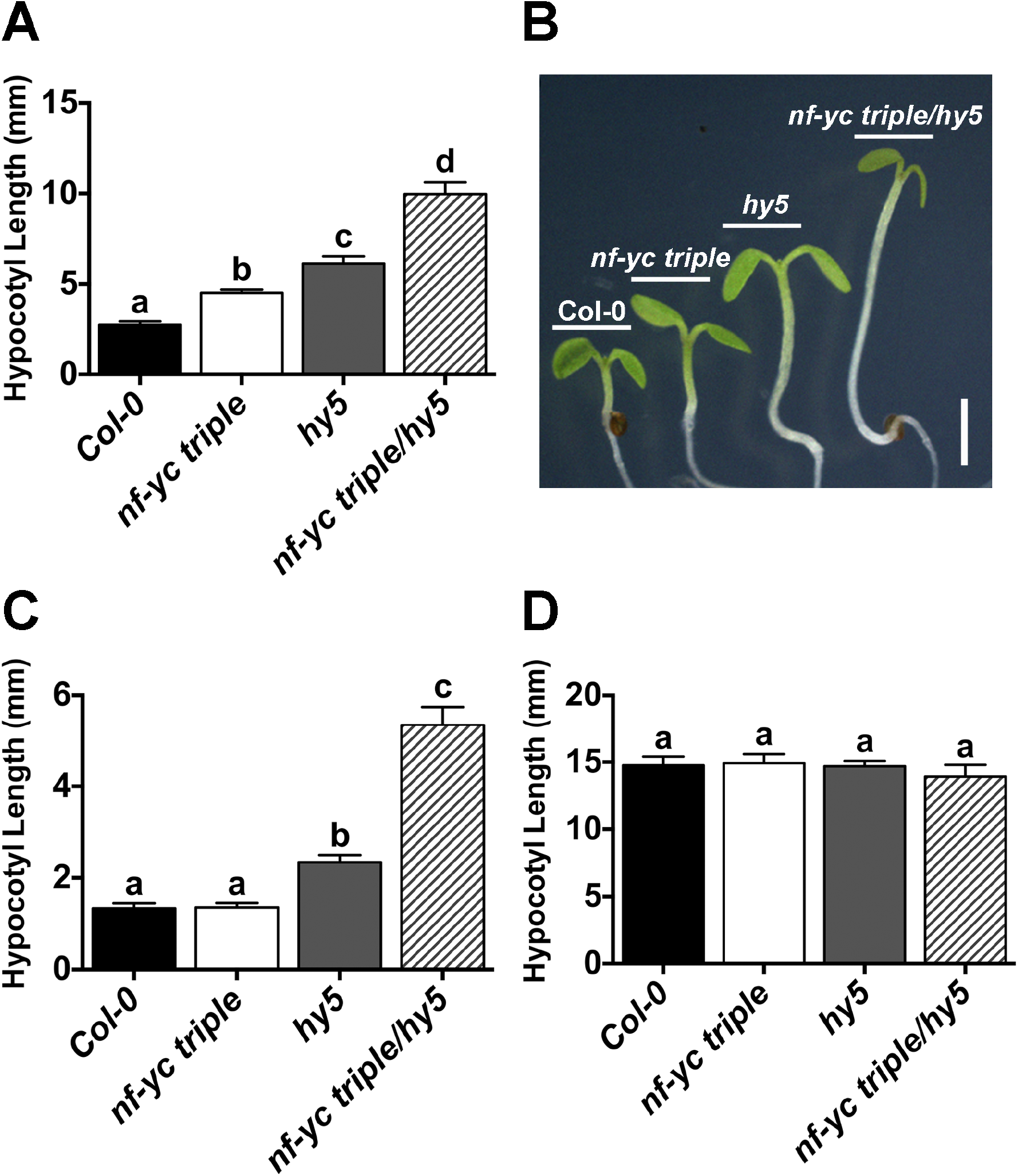
Light perception is synergistically defective in *nf-yc triple hy5* mutants. Hypocotyl lengths are shown for five day old plants grown on B5 media in **A-B**) SD, **C**) cWL, and **D**) cD. Statistically significant differences were determined and described in Figure 2. Scale bar in B) represents 2mm.

### NF-YC3, 4, and 9 and HY5 have both shared and independent regulatory targets

To further dissect the genetic relationship between NF-YC and HY5, we compared the transcriptome profiles of seven day old, cWL grown *nf-yc triple, hy5*, and *nf-yc triple hy5* mutant seedlings using RNA Sequencing (RNASeq).

When compared to wild type, *hy5* mutants had 1368 up-regulated and 941 down-regulated genes, whereas analysis of differentially expressed genes in the *nf-yc triple* mutant showed a smaller set of 645 up-regulated genes and 493 down-regulated genes (at least 1.5 fold, adjusted *p* < 0.05, Table S1). Direct comparison of the *hy5* and *nf-yc triple* down-regulated genes showed substantial overlap, with approximately 40% of the *nf-yc triple* down-regulated genes being contained in the *hy5* data set (Fig 5A). Geneontology (GO) analysis for genes down-regulated in both the *nf-yc triple* and *hy5* mutants identified enrichment in many categories involved in photomorphogenesis and early seedling development, including response to light stimulus and pigment biosynthetic processes (Table S2). Comparison between up-regulated gene sets yielded similar results with ~50% shared between the *nf-yc triple* and *hy5* (Fig 5B). GO enrichment analyses of genes up-regulated in both *nf-yc triple* and *hy5* yielded categories in cellular stress responses and cellular responses to hormones, including ethylene, salicylic acid, and jasmonic acid (Table S3).

**Figure 5.**
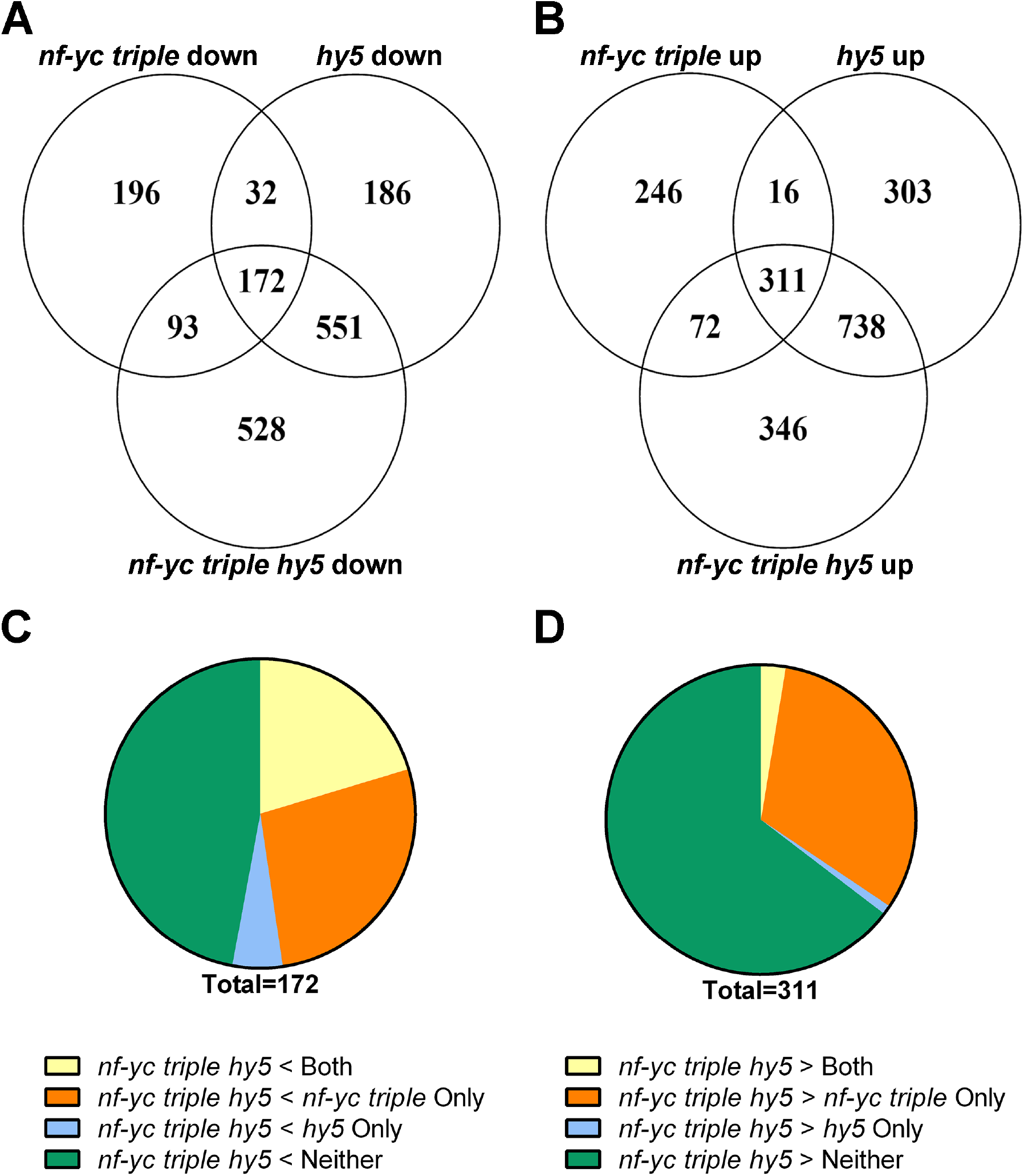
RNASeq analysis identifies both shared and independently regulated targets for NF-YCs and HY5. Overlap between genes significantly **A**) down-regulated or **B**) up-regulated at least 1.5-fold in the *nf-yc triple, hy5*, and *nf-yc triple hy5* backgrounds, relative to Col-0. Genes significantly misregulated in all three genotypes were then broken into regulatory groups according to the level of **C**) down-regulation or **D**) up-regulation in the *nf-yc triple hy5* mutant relative to both *nf-yc triple* and *hy5*.

To further investigate the regulatory relationship between HY5 and the NF-YCs, we analyzed genes that were either significantly up-regulated or down-regulated in *nf-yc triple hy5* mutants (Table S1). These data sets were then sub-divided into four groups based on the level of misregulation (defined as either significantly up-regulated or down-regulated) in the *nf-yc triple hy5* mutant relative to both *nf-yc triple* and *hy5:* Group I) Genes more misregulated in the *nf-yc triple hy5* mutant than both *nf-yc triple* and *hy5*; Groups II-III) Genes more misregulated in the *nf-yc triple hy5* mutant compared only to the *nf-yc triple* (II) or *hy5* (III); and Group IV) Genes not misregulated compared to either the *nf-yc triple* and *hy5* (i.e., still misregulated in the quadruple mutant, but no change from *nf-yc triple* and *hy5* (Fig 5C–D)). GO enrichment analyses of these four groups represent putative biological processes that NF-YCs and HY5 regulate cooperatively (genes more misregulated in *nf-yc triple hy5* than parental lines) and independently (genes not misregulated in *nf-yc triple hy5* relative to *nf-yc triple* and/or *hy5* (Tables S2–3)).

Analysis of genes significantly more down-regulated in the *nf-yc triple hy5* mutant (relative to its parental genotypes) identified several over-represented categories, including flavonoid biosynthesis and polyol metabolic processes (Table S2). Among genes up-regulated to a similar level in the *nf-yc triple hy5, nf-yc triple*, and *hy5* data sets, and consistent with the synergistic hypocotyl phenotype of the *nf-yc triple hy5* mutant, there was a significant enrichment for genes involved in cell wall organization, cell wall biogenesis, and cell wall macromolecule metabolic processes (Table S3). Taken together, these data identify putative targets and biological processes regulated both cooperatively and independently by NF-YCs and HY5, solidifying the existence of a complex functional relationship.

### Longer hypocotyls in nf-yc triple hy5 mutants is largely a function of greatly increased cell elongation

Previous research established that the elongated hypocotyls in *hy5* mutants are directly related to increased epidermal cell length (18); therefore, we additionally examined individual files of epidermal cells along the hypocotyls of *nf-yc triple hy5* mutants for total cell number and mean cell length (Fig 6A–C). The mean length of individual epidermal cells in *hy5* (82 μm) was ~90% greater than Col-0 (43 μm), while cells in the *nf-yc triple* mutant measured only ~15% longer than Col-0. Reflecting the synergistic hypocotyl elongation phenotypes of *nf-yc triple hy5* mutants, the epidermal cells of the quadruple mutant (158μm) were ~270% longer than those measured in Col-0. Total epidermal cells in the quadruple mutant were also increased >60% compared to Col-0. Therefore, the very long hypocotyls of *nf-yc triple hy5* mutants can be explained by a combination of modestly increased cell count and highly increased cell elongation.

**Figure 6.**
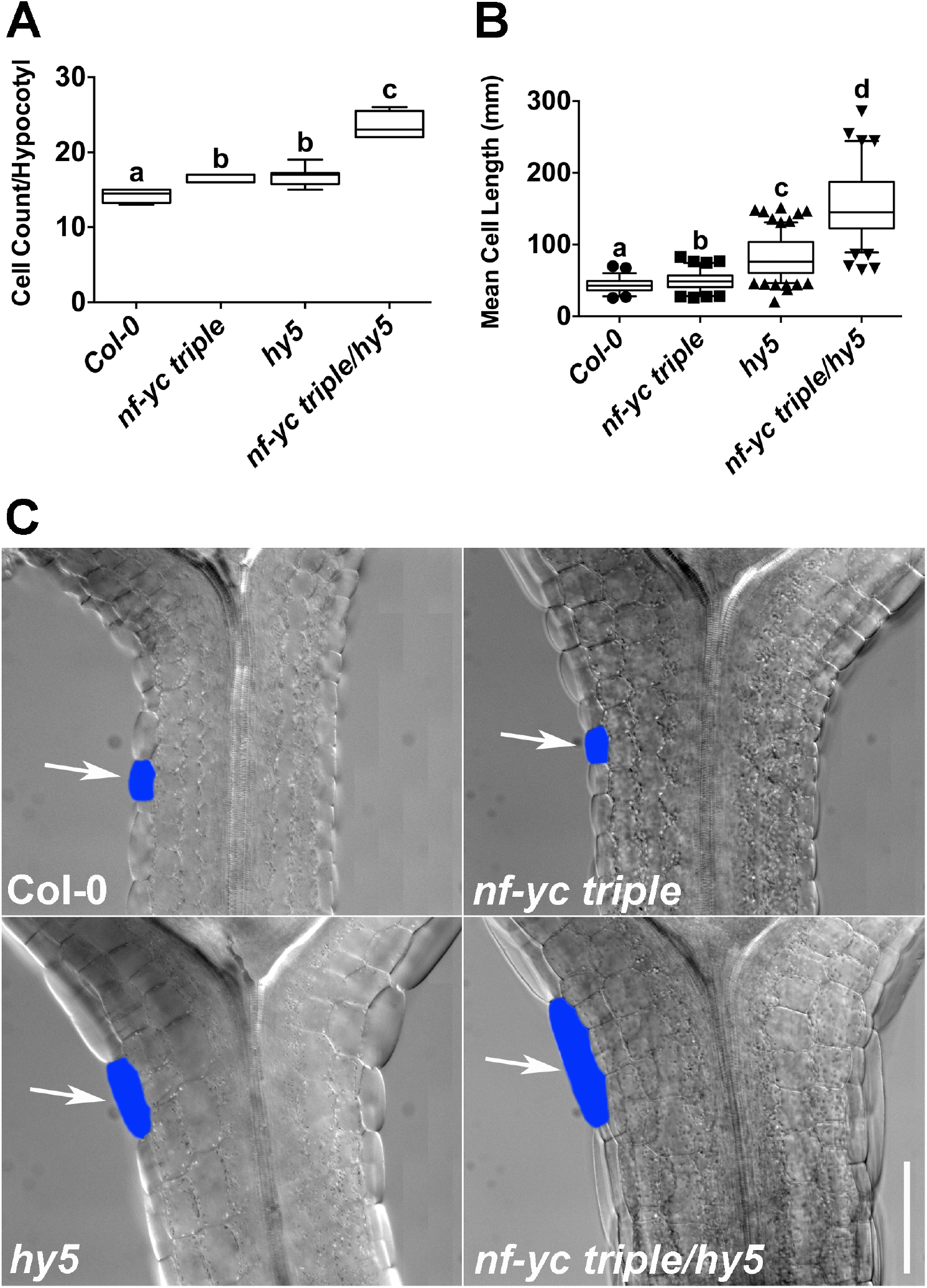
Synergistically longer hypocotyls in *nf-yc triple hy5* mutants are a function of moderately more cells and greatly increased cell elongation. Single linear files of hypocotyl epidermal cells were both **A**) counted and **B**) measured for mean length. Plant were grown in cWL and measurements were taken on five day old plants. **C**) Example hypocotyls for each genotype near the cotyledon junction – blue color marks a representative single cell in each genotype. Arrows point to typical epidermal cells for each genotype. Scale bar = 100μm.

### NF-YC transcription factors suppress hypocotyl elongation via pathways at least partially independent of HY5, HFR1, and LAF1

HY5 regulates photomorphogenesis regardless of wavelength, whereas HFR1 and LAF1 are more specific to FR light responses (24–28). To better compare the spectrum of *nf-yc* mutant defects to these other transcription factors, we first examined both the *nf-yc triple* and *nf-yc triple hy5* lines in cB, cFR, and cR over a gradient of light intensities (Fig 7A–C). Under all but the lowest cB fluence rates, the *nf-yc triple hy5* mutants had significantly longer hypocotyls than all other lines (Fig 7A). Considering our previous observation that *nf-yc triple* light perception defects were only apparent in SD conditions (Fig 1B), it was somewhat surprising to find that *nf-yc triple hy5* defects in cB were most pronounced at the highest light intensities (Fig 7A).

**Figure 7.**
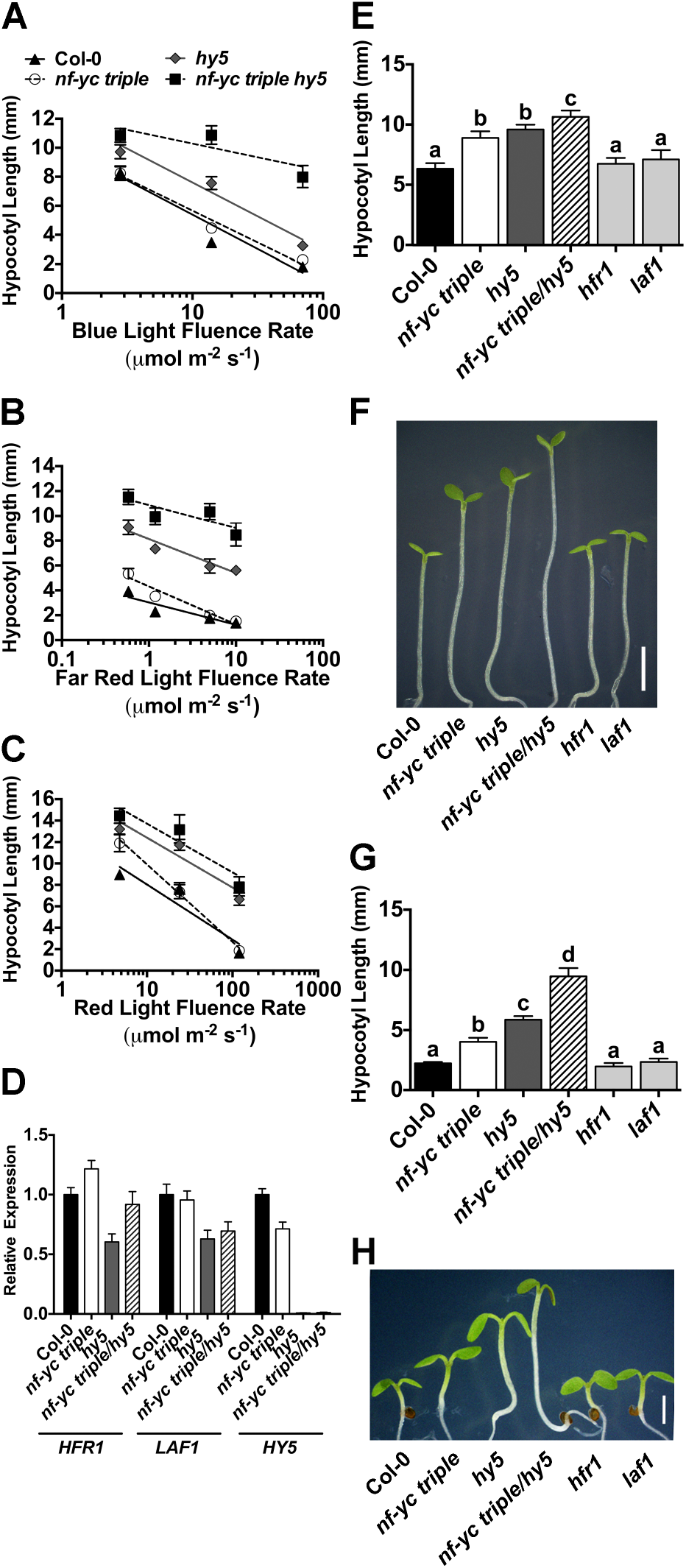
*nf-yc* mutants have broad light perception defects that are at least partly independent of *HFR1, HY5*, and *LAF1*. Fluence rate curves for hypocotyl lengths are shown for five day old plants in **A**) cB, **B**) cFR, and **C**) cR light conditions (see symbols key in A). **D**) qPCR of *HFR1, LAF1,and HY5* in key genetic backgrounds. **E-F**) Quantification and images of typical phenotypes for mutants grown in low fluence rate cR (4.8μmol m^−2^ s^−1^). Scale bar in F = 2mm. **G-H**) Quantification and images of typical phenotypes for mutants grown in SD, white light conditions. Scale bar in H = 2mm. Statistically significant differences were determined as described in Figure 2.

The *nf-yc triple hy5* mutants were significantly longer than their *nf-yc triple* and *hy5* parental lines under all cFR conditions (Fig 7B). One possible cause for defects in FR light perception could be misregulation of *HY5, HFR1*, or *LAF1* in *nf-yc* mutants – i.e., NF-Y complexes could control the expression of these genes. Figure S2 shows that *HY5* is not misregulated in an *nf-yc triple* background in cWL and we additionally examined the expression of *HY5, HFR1*, and *LAF1* in cFR grown plants. Consistent with previous reports (35, 74), modest differences in *HY5, HFR1, and LAF1* were either insignificant or not reproducible in repeated expression analyses in the various mutant backgrounds (Fig 7D). We conclude that *nf-yc* mutant phenotypes are not likely related to simple changes in the expression of these well-known regulators of light perception. Further, as discussed below, *nf-yc triple* mutants appear to have a different spectrum of light defects than either *hfr1* or *laf1* mutants.

Under low fluence rate cR and cFR, the *nf-yc triple* mutant alone had significantly longer hypocotyls than Col-0, which is similar to *hy5* (Fig 7B–C). However, *hfr1* and *laf1* are only reported to have defects in cFR (both) and cB light (*hfr1*). To confirm these previous reports with our experimental conditions, we directly compared *hfr1* and *laf1* to *nf-y* mutants under low fluence rate cR (Fig 7E–F). As previously reported, *hfr1* and *laf1* appeared identical to wild type Col-0 plants, whereas the *nf-yc triple* mutants were consistently ~40% longer than Col-0 and similar to *hy5* mutants. We additionally compared the *nf-yc triple* to *hfr1* and *laf1* in SD conditions (Fig 7G–H). As expected, the *hfr1* and *laf1* mutants appeared phenotypically identical to Col-0, while the *nf-yc triple, hy5*, and *nf-yc triple hy5* mutants were all significantly longer. Collectively, our data suggests that NF-YCs regulate hypocotyl elongation via an independent pathway(s) from HY5, and at least partially independent of HFR1 and LAF1, with broad roles in light perception at variable fluence rates.

### Loss of NF-YC function partially suppresses cop1 mutant phenotypes

In the dark, HY5, HFR1, and LAF1 are all targeted for degradation by the proteasome in a COP1-dependent manner (28, 75, 76). COP1 mutants have short hypocotyls and other photomorphogenic phenotypes even when grown in the dark, and these phenotypes are partially suppressed in *hy5 cop1, laf1 cop1*, and *hfr1 cop1* double mutants (75, 77, 78). Therefore, we examined if the *nf-yc triple* mutation could also suppress the short phenotype of dark-grown *cop1-4* mutants. Similar to *cop1 hy5*, an *nf-yc triple cop1* mutant had ~80% longer hypocotyls than the *cop1* single mutant when grown in constant darkness (Fig 8A). Because *cop1-4* mutants are known to have reduced rosette diameters (dwarf phenotype) and early flowering (79), we further characterized these phenotypes in *nf-yc triple cop1* mutants. For rosette diameter, the *nf-yc triple* mutant was once again able to partially suppress *cop1* (Fig 8B). One possibility is that this suppression is simply a function of *nf-yc triple* mutants being late flowering - i.e., because they are later flowering, the rosettes have time to achieve a greater diameter prior to the phase change to reproductive growth. However, a control cross between *cop1* and an even later flowering *constans* mutant (the alternatively named *co-sail* or *co-9* allele (80)) had no impact on rosette diameter when crossed to *cop1-4*. This suggests that *nf-yc* loss of function alleles genuinely suppress the small *cop1* rosette diameter phenotype and that this NF-YC function is genetically separable from its role in flowering time.

**Figure 8.**
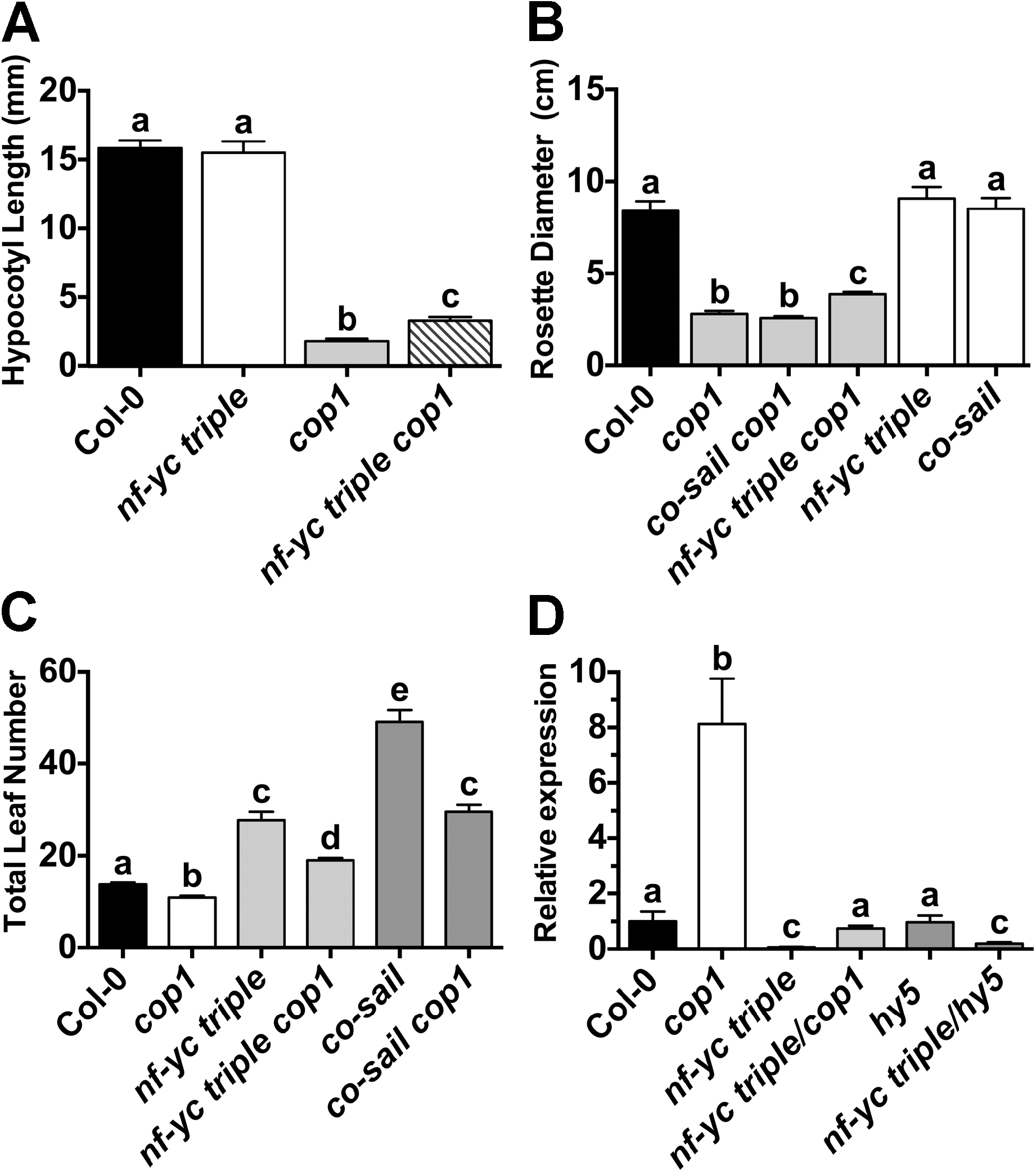
Multiple *cop1-4* mutant phenotypes are partially dependent on *NF-YC* genes. Partial suppression of cop1 mutant phenotypes are quantitated for **A**) dark-grown seedling hypocotyl elongation, **B**) rosette diameter, **C**) flowering time, and **D**) relative *FT* expression levels. Statistics and labeling as in Figure 2, except *FT* expression statistics which were calculated using qBase software (Biogazelle).

Finally, we also tested whether the *nf-yc triple* mutant could suppress the early flowering phenotype of *cop1* (Fig 8C). The *nf-yc triple cop1* mutant plants flowered moderately, but significantly, later than Col-0, intermediate to the early flowering *cop1-4* mutant and late flowering *nf-yc triple* mutant. This result is not surprising as an important role for COP1 in flowering is to suppress CO function via protein degradation (79, 81). Because CO and NF-Y function together to regulate photoperiod-dependent flowering (49, 52, 57, 82), the basis of early flowering in *cop1* is largely a function of CO protein (and potentially NF-Y, see below) hyper-accumulation (79, 81). Measurements of *FLOWERING LOCUS T (FT*) expression – the regulatory target of CO and NF-Y function in flowering time – correlated with expectations from the flowering time measurements (Fig 8D).

### Like HY5, HFR1, and LAF1, NF-YA proteins are degraded in the dark

NF-YC regulation of light perception appears to share many parallels with HY5, HFR1, and LAF1, including the suppression of *cop1* mutant phenotypes (75, 77, 78). As with these other photomorphogenic transcription factors, it is tempting to speculate that NF-YC proteins might be targets of COP1-mediated proteasome degradation in the dark. However, this does not appear to be the case as native antibodies to both NF-YC3 and NF-YC4 show modest fluctuations, but largely stable accumulation throughout both short day and long day cycles (Fig S5 – recall also that expression of any *one* NF-YC from a native promoter rescues the mutant phenotype, Fig S3). Nevertheless, NF-YC proteins function within the context of a heterotrimeric complex and reduction of the NF-YA or NF-YB components could also disrupt activity.

In this regard, overexpression of most NF-YAs leads to small, dwarf phenotypes that are not unlike those observed for *cop1* mutants (58, 83). In fact, when we examined *NF-YA* overexpressing plants (35S promoter driven; previously described in (58)), they were found to have significantly shortened hypocotyls in both cD and cR conditions (Fig 9A). While shortened hypocotyls in cD is a classic constitutive photomorphogenic phenotype, expressing *p35S:NF-YA* hypocotyl lengths in cR as a percentage of cD growth additionally showed that most of these plants were specifically defective in red light perception (Fig 9A).

**Figure 9.**
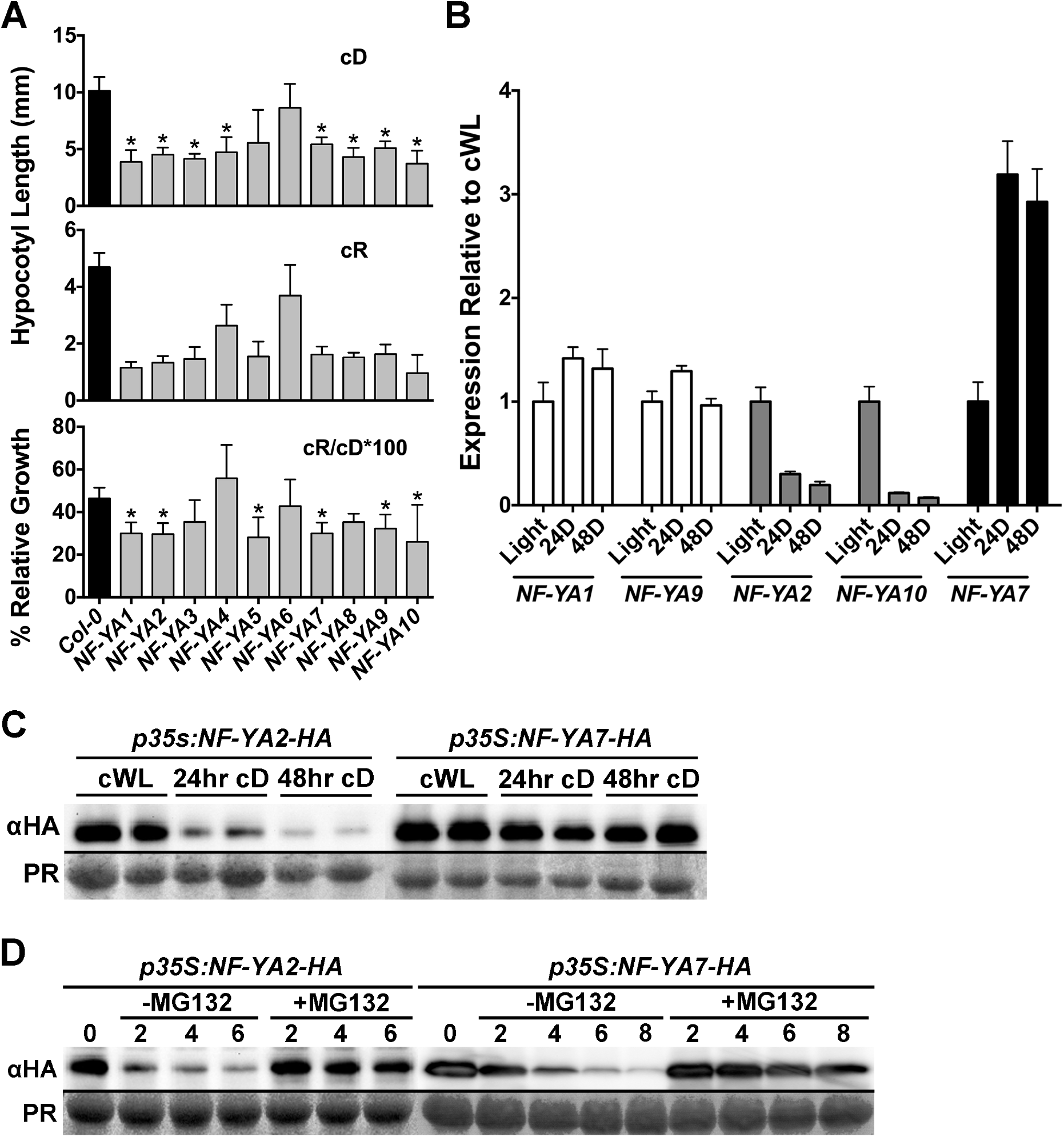
NF-YAs represent the light/dark cycle regulated components of NF-Y complexes. **A**) *NF-YA* overexpression causes constitutive photomorphogenesis phenotypes as well as cR light perception defects. To account for the constitutive photomorphogenic phenotypes of *NF-YA* overexpressing plants, relative percent growth in cR was determined by dividing cR hypocotyl length by cD hypocotyl length and multiplying by 100. **NOTE**: Because of their cop1-like, dwarf phenotypes, seed numbers are limited for NF-YA overexpressing plant. Thus, we chose to focus on cR light perception because of the relatively strong light defects measured for the *nf-yc triple* (Figure 2C). Statistically significant differences were determined as described in Figure 2. Error bars 95%. **B**) Expression of select *NF-YA* genes after 24–48hrs darkness relative to cWL. Error bars SEM. **C**) Protein accumulation of 35S promoter-driven *NF-YA2* and *NF-YA7* constructs in cWL compared to 24–48hrs cD. **D**) Cell free protein degradation assays for NF-YA2 and NF-YA7, with and without the proteasome specific inhibitor MG132.

While it is unknown which of the 10 Arabidopsis NF-YAs is natively involved in hypocotyl elongation, a recent publication suggested that NF-YA2 may be found in complex with NF-YC3, 4, and 9 (84). Therefore, using qPCR, we examined the expression of *NF-YA2* in 24hr cWL or after 24–48hrs of cD and found that expression was strongly downregulated in cD (Fig 9B). At the same time we compared *NF-YA2* expression to a subset of other *NF-YA* genes - *NF-YA1*, 7, *9*, and *10. NF-YA10* is the most closely related paralog to *NF-YA2* (encoding 63% identical full length proteins, (85)) and it showed the same pattern of downregulation in cD. However, the less related *NF-YA1* and *9* genes (42% identical to each other, but only 23–24% to NF-YA2) remained stably expressed in cD, while *NF-YA7* was actually upregulated. Thus, expression of the *NF-YA* gene family in response to cD is quite variable, and suggests potential for light regulated accumulation and depletion.

To determine if NF-YA proteins might be targets of degradation in the dark, we examined the accumulation of NF-YA2 and NF-YA7 expressed from constitutive 35S promoters. We chose to use a constitutive promoter to differentiate between changes in protein accumulation due to reduced gene expression (see Fig 9B) versus active protein degradation processes. Strongly suggesting an active degradation process, NF-YA2 protein accumulation was strongly reduced in cD conditions, even when expression was driven from the 35S promoter (Fig 9C). This was in stark contrast to NF-YA7 which had stable protein accumulation in the dark. To determine if the proteasome was involved in the process, we additionally performed cell-free protein degradation assays (as previously described (86)) and determined that NF-YA2 was rapidly degraded (Fig 9D). However, the addition of the proteasome inhibitor MG132 strongly reduced the apparent degradation of NF-YA2 protein. We note that NF-YA7 also degraded in an MG132 dependent manner, suggesting that it can also be targeted by the proteasome for degradation, even if darkness may not be the driving force (Figs 9B–C). Collectively, these data suggest that NF-YAs can also regulate light perception and are targeted for proteasome mediated degradation, perhaps controlling the overall stability of the NF-Y complexes necessary to suppress hypocotyl elongation in the light.

## Discussion

The physical and genetic relationships between the NF-YCs and HY5 are similar to those reported for several other photomorphogenic transcription factors, such as HFR1 and LAF1, where both are able to physically interact with HY5, but have clearly additive light perception defects when combined in higher order mutants (24, 35, 77, 81). While we observed *nf-yc triple* mutant phenotypes primarily in low-intensity monochromatic light, the most pronounced *nf-yc triple hy5* phenotypes (relative to both the *nf-yc triple* and *hy5* mutants) were seen in high-intensity monochromatic light. Because HY5 protein accumulation and activity are light-intensity dependent (23), one possible explanation for this relationship, supported by our RNASeq data, is that HY5 and the NF-Y complexes share a subset of regulatory functions. When *nf-yc triple* mutants are grown in low light, HY5 function becomes essential for maintaining normal photomorphogenesis; however, under these same conditions, HY5 is also increasingly targeted for degradation and the nf-y-related photomorphogenic defect becomes apparent. Alternatively, in saturating, high light conditions, the relative NF-Y contribution to photomorphogenesis is masked due to high accumulation and activity of HY5 and other photomorphogenic transcription factors.

### Are NF-YC by HY5 Interactions Important in Light Signaling?

Unexpectedly, while we were able to detect a physical interaction between NF-YC9 and HY5 through FRET-FLIM analyses, we were not able to detect an interaction between NF-YC3 or NF-YC4 and HY5. This is surprising because the histone fold domains of NF-YC3, 4, and 9 are nearly identical (in fact, NF-YC3 and NF-YC9 are identical (85)); however, the amino- and carboxy-terminal regions are more divergent and could be involved in the NF-YC by HY5 physical interaction. Because of the extreme spatial constraints required for FRET to occur, it is not valid to conclude from these experiments that NF-YC3 and NF-YC4 cannot interact with HY5 (87, 88), and Y2H analyses did previously show a positive interaction between NF-YC4 and HY5 (57). The question also remains whether or not the ability of NF-YCs to physically interact with HY5 is of biological importance relative to their specific functions in light signaling. Both HY5 and the NF-YCs are also regulators of ABA signaling (19, 57, 58) and it is possible that a physical interaction between them is only related to this or another undefined pathway. This possibility is supported by the additive, and even synergistic, mutant phenotypes of the *nf-yc triple hy5* plants – i.e., if these proteins are physically interacting in a linear pathway or at a common hub in light signaling, how does the quadruple *nf-yc3 nf-yc4 nf-yc9 hy5* mutation result in these synergistic phenotypes? Alternatively, arguing for the relevance of physical interactions in light signaling, there is clearly a significant amount of overlap in the putative regulatory targets of NF-YC3, 4, 9 and HY5. Similarly, it was also previously suggested that subsets of photomorphogenic responses might be regulated by combinations of both overlapping (where physical interactions were relevant) and non-overlapping functions between HY5, HFR1, and LAF1 (35). In future experiments, it will be informative to examine the stability of NF-YA proteins in the presence or absence of these other transcriptional regulators as it was previously shown that HFR1 and LAF1 were co-dependent for their protein stability (i.e., they required each other to avoid proteasome-mediated degradation; (74)). Ultimately, deciphering these putative cooperative versus individual roles remains an exciting challenge for future research.

### Is there a separate pathway for NF-YC function in light perception?

While we have demonstrated that NF-YC3, 4, and 9 function at least partially independently of HY5 in light perception, one pressing question is whether the NF-Y complex is functioning through other known light-responsive transcription factors, such as HFR1 or LAF1. Directly addressing this hypothesis would require the creation of *nf-yc triple hfr1* and *nf-yc triple laf1* quadruple mutants; however, because *HFR1* is linked to *NF-YC9*, traditional crossing techniques would be prohibitively difficult. Therefore targeting of these loci in the *nf-yc triple* mutant with CRISPR/Cas is currently underway. To support that the NF-YCs are functioning separately, or at least differently, from HFR1 or LAF1, we identified strong phenotypes in the *nf-yc triple* mutant in SD- and low cR-grown seedlings, where *hfr1* and *laf1* showed no mutant phenotype. Additionally, we found no misregulation of *HFR1* and *LAF1* in cFR-grown *nf-yc triple* mutants. These data suggest that the function of NF-YCs in light perception is at least partially separable from the function of HFR1 and LAF1.

Similar to the NF-YCs, both STH2/BBX21 and STH3/BBX22 function as photomorphogenesis-activating transcription factors over a broad range of light conditions and are also able to physically interact with HY5 (29, 30). It is possible that the NF-YCs are functioning in an STH2/STH3-dependent manner; however, phenotypes of *sth2 sth3* double mutants and *sth2 sth3 hy5* triple mutants suggest that this might not be the case. In contrast to the genetic relationship between the *nf-yc triple* and *hy5*, the hypocotyls of *sth2 sth3 hy5* triple mutants were not longer than *hy5* in cR. Further, we observed the most severe *nf-yc triple* phenotypes in low-intensity light while the *sth2 sth3* mutant phenotypes were only observed in high-intensity light (30). Nevertheless, these observations do not preclude genetic interactions for subsets of shared functions, similar to what we have already suggested with HY5.

When examining the protein domains of STH2/BBX21 and STH3/BBX22, it is tempting to speculate that there may be indirect physical interactions with the NF-YC proteins as part of a larger light perception complex. STH2 and STH3 have B-box domains, thus their alternate BBX21 and BBX22 designations (89), and these domains are necessary for direct physical interactions with HY5 (29, 30). BBX proteins also often have so-called CO, CO-LIKE, and TIMING OF CAB (CCT) domains (89, 90). For example, CO (BBX1) is a BBX-CCT protein and mutations in either of these domains impacts its ability to regulate flowering time (91). It is well-established that NF-YC3, 4, and 9 can all physically interact with CO and the CCT domain is both necessary and sufficient for this interaction (49, 51, 52). While STH2 and STH3 do not have a CCT domain, recent evidence demonstrated that BBX proteins can heterodimerize with other BBX proteins (92). Therefore, it is possible that NF-Y complexes may interact with BBX-CCT proteins via the CCT domain and recruit other non-CCT containing BBX proteins, such as STH2 and 3, to these complexes.

### Are NF-Y complexes activators of photomorphogenesis or skotomorphogenesis? Or both?

In contrast to our genetic evidence showing that NF-Ys act as suppressors of hypocotyl elongation, NF-YB9/LEC1 appears to have the opposite role. This idea comes from recent evidence demonstrating that inducible overexpression of *NF-YB9/LEC1* also resulted in elongated hypocotyls, suggesting that NF-YB9/LEC1 might actually function as an enhancer of hypocotyl elongation (38, 93). Consistent with this finding, embryonic hypocotyls are shortened in *lec1* mutants (62). Further, recent data shows that the hypocotyls of both light and dark grown *lec1* mutants are significantly shorter than wild type plants (39). Interestingly, overexpression of a repressor of photomorphogenesis - PHYTOCHROME-INTERACTING FACTOR 4 - results in elongated hypocotyls, but this phenotype is partially dependent on the presence of *LEC1* (39). These results raise a few interesting questions: why would NF-YB9 act opposite to the NF-YA and NF-YC members of the complex (as reported here) and what mechanism would allow this result? Considering these questions, it is important to remember that each NF-Y subunit – A, B, and C – is part of a 10 member family (41). Thus, many unique NF-Y complexes could theoretically form and, depending on their composition, actually act to competitively suppress or enhance a given process. In this scenario, some members of a given NF-Y family might enter a complex, but render it inactive, while other members of the same family would have the opposite effect.

Our previous research on ABA-mediated seed germination provides some precedence for the above idea. We demonstrated that members of the same NF-Y family can act in opposing manners, either enhancing or delaying germination when their expression is altered (both NF-YA and NF-YC examples exist; (57, 58)). Similarly, some BBX proteins also show these opposing functionalities. For example, BBX24 and 25 are hypothesized to interfere with BBX22 function by entering into non-functional complexes with HY5 (94). Fitting this scenario nicely, NF-YB9/LEC1, and its closest relative NF-YB6/LEC1-LIKE, are quite unique and very different from the other eight NF-YB proteins in Arabidopsis. This includes 16 amino acid differences in their highly conserved histone fold domains that are completely unique to only this pair (95). However, an alternative hypothesis must be considered related to the most recent *lec1* data (39): *lec1* shortened hypocotyls may not be developmental patterns related to loss of skotomorphogenesis or post embryonic in nature, but, instead, are lasting patterns laid down during embryogenesis. This possibility is supported by both the modest magnitude of the effects and the finding that *lec1* plants are short in all conditions (dark and light). This is not the case for the *nf-yc* mutants reported here as they are indistinguishable from wild-type plants in the dark and elongated in the light, clearly defining them as positive regulators of photomorphogenesis.

### NF-Y complexes share the hallmarks of photomorphogenic transcription factors

While the functional relationship between the NF-YCs and HY5 is similar to that observed with many other photomorphogenic transcription factors, the NF-YCs do not appear to be transcriptionally or translationally regulated in a manner consistent with light-responsive proteins; however, because the NF-YCs act in the larger context of an NF-Y trimer, the physical properties and regulatory components of the functional unit can be spread across multiple proteins. We showed that several NF-YA subunits with photomorphogenic phenotypes are regulated by light, and that NF-YA2 is targeted for degradation by the proteasome. Regulation of the NF-YA subunit establishes NF-Y complexes as possessing all of the properties expected of a photomorphogenic transcription factors, including DNA-binding capacity, the ability to physically interact with other photomorphogenic factors, and a light-regulated mechanism to modulate function and abundance. While specific NF-YA subunits have not been identified to natively regulate the inhibition of hypocotyl elongation, a recent publication showed that over-expression of NF-YA2 led to earlier flowering (84). This suggests that NF-YA2 could be integrated into an NF-Y complex containing NF-YC3, 4, and/or 9, as each is redundantly involved in photoperiod-dependent flowering (51); finally, an NF-YA2/NF-YB2,3/NF-YC9 trimer has been identified through yeast three-hybrid analyses, and further verified through two-way interaction assays (including BiFC and co-IP, (84)). The identity of photomorphogenic NF-YB proteins remains unknown and it will interesting to determine which, if any, non-LEC-type NF-YB will be positive regulators of photomorphogenesis.

## Conclusion

The data presented here firmly establishes NF-Y complexes as positive regulators of photomorphogenesis, significantly extending recent findings (38, 39). Future research on the potential regulation of NF-YA proteins by the proteasome and the identity of photomorphogenic NF-YA and NF-YB will improve our current understanding. Although only discussed at a cursory level here, research on NF-Y roles in flowering time demonstrate important interactions with the BBX protein CONSTANS and suggest that NF-Y by BBX interactions may be generalizable (49–52, 96). Given the numerous roles for BBX proteins in light perception (29, 30, 94, 97–99), we predict that future studies will uncover BBX by NF-Y interactions that are essential for light perception. This would be an exciting finding, significantly extending the regulatory reach and capacity of the four interacting families of proteins.

## Methods

### Growth Conditions and Plant Lines

All plants were of the Col-0 ecotype and were grown at 22°C. Prior to starting germination on plates or soil, seeds were cold-stratified in a 4°C walk-in cooler in the dark for 2–3 days. Plants grown in continuous white light were grown in a Conviron ATC13 growth chambers or a custom walk-in growth chamber. Plants in single wavelengths light experiments were grown in a Percival E30-LED growth chamber after initial exposures to 4 hours of white light to induce germination. Plants used in flowering-time experiments, rosette diameter measurements, and GUS staining were grown in a previously-described soil mixture (NF-YC Triple Flowering Time Paper Ref). All other plants were grown on 0.8–2% agar plates supplemented with Gamborg B-5 Basal Medium (PhytoTechnology Laboratories, product #G398). *Nicotiana benthamiana* plants were grown under long-day conditions (16h light/8h dark) at 22°C in a Conviron ATC13 growth chamber. For UV experiments, plants were grown in cWL for 5 days under a Mylar filter (Professional Plastics, catalog #A736990500) or mock-filter. Mutant lines used in this study, including references for their original derivation and description in the literature, are reported in Table S4 (19, 24, 51, 100–104). AGI identifiers for all genes reported are also described in Table S4.

GUS Staining, Rosette Diameter Measurements, and Flowering Time Experiments GUS staining was performed as previously described on 5 day old soil-grown seedlings (105), and images were taken on a Leica dissecting stereoscope. Flowering time was measured as the total number of rosette and cauline leaves present shortly after bolting (106), and all genotypes exhibited similar developmental rates. To quantify rosette diameter, plants were photographed from above at the time of bolting, and the Feret’s diameter was measured, anchored at the tip of the longest rosette leaf.

### Hypocotyl Length and Cell Length Measurements

To measure hypocotyl elongation, seeds were sown onto B5 supplemented plates with 2% agar and cold-stratified at 4°C for 2 days. Before transfer to specific light conditions, all plates were set at room temperature in continuous white light for 4 hours. Plates were grown vertically for the duration of the experiments. Germination rates for the *nf-y* mutants under study were previously shown to be the same in B5 media and confirmed for the experiments reported here (57). To facilitate proper measurement, all plants were straightened on the plate before taking pictures. Pictures were processed, and all individual hypocotyls were traced and measured in FIJI (107).

For individual cell length and total cell number measurements of single files of hypocotyl cells, seedlings were grown on plates as described above for 5 days in cWL. Seedlings were fixed in an FAA solution and dehydrated through sequential 30-minute incubations in 90% and 100% ethanol (108). Fixed and dehydrated seedlings were individually mounted in the clearing agent methyl salicylate (108), and immediately taken for measurement on a Nikon Eclipse NI-U compound microscope. Using Differential Interference Contrast (DIC) optics, individual cell files were identified and measured manually through the NIS-Elements BR software, and pictures were taken at the hypocotyl-cotyledon junction for every seedling measured.

### qRT-PCR Analyses

Total RNA was isolated from 7-day-old seedlings grown under continuous white light conditions and from 5-day-old seedlings grown under continuous far red light conditions using the Omega Biotek E.Z.N.A Plant RNA Kit (catalog #R6827–01), and was DNase treated on-column with Omega Biotek’s RNase-free DNase set (catalog #E1091). First-strand cDNA synthesis was carried out with Invitrogen’s SuperScript^®^ III Reverse Transcriptase (catalog #18080–044) and supplied oligo dT primers. qRT-PCR was performed on a Bio-Rad CFX Connect Real-Time PCR Detection System (http://www.bio-rad.com/), using Thermo Scientific’s Maxima SYBR Green/ROX qPCR Master Mix (catalog #K0222). Each genotype was assayed with three independent biological replicates, consisting of approximately 100mg of starting tissue each. White light grown seedlings were normalized to *At2g32170*, while far red light grown seedlings were normalized to *At3g18780* and *At1g49240*. Statistical analysis and comparisons between samples was performed in the Bio-Rad CFX Manager Software (http://www.bio-rad.com/) through use of the 2^(−ΔΔCT)^ method.

### Transient Transformation of *Nicotiana benthamiana* Leaves

The leaves of 4- to 6-week old *N. benthamiana* were co-infiltrated with *Agrobacterium tumefaciens* GV3101 strains harboring either a YFP-fused protein or an mCer3-fused protein, in addition to the *Agrobacterium* strain C58C1 harboring the viral silencing suppressor helper complex pCH32 (109). Before infiltration into *Nicotiana* leaves, *Agrobacterium* cultures grown overnight were treated with 200uM acetosyringone in a modified induction buffer for 4 hours (110). This induced culture was re-suspended in 10mM MES 10mM MgSO4 and directly infiltrated into young leaves. All downstream analyses were conducted 2–4 days after initial infiltration.

### FRET-FLIM and FRAP Analyses

FLIM data was acquired through time-correlated single photon counting (TCSPC) on a Lecia TCS SP8 confocal laser scanning microscope using an HC PL APO 40x/1.10 water immersion objective. Fluorescent protein excitation was achieved through use of a titanium-sapphire multiphoton laser (Chameleon, Coherent) operating at 120 femtosecond pulses of 858nm infrared light. Fluorescence emissions were detected by non-descanned hybrid detectors (HyDs). Fluorescence lifetimes of entire nuclei were fit to a single-exponential model through the SymphoTime 64 (www.picoquant.com) software, and comparison of the fluorescence lifetimes before and after FRAP was used to detect FRET. For FRAP analyses, YFP photobleaching was accomplished with a high-intensity Argon laser line at 514nm for 15 seconds, followed by recovery imaging of both mCer3 (excited at 458nm) and YFP (excited at 514nm) every second for 5 seconds. Descanned HyDs were used to detect mCer3 emission from 459nm to 512nm, and a Photomultiplier Tube (PMT) was used to detect YFP emission from 512 to 562nm, with a 458/514 notch filter in place. This process was performed a total of 3 times for each nucleus, and both mCer3 and YFP intensities were calculated relative to initial fluorescence intensity. FRAP was conducted as an internal control during FLIM measurements, allowing us to assess the level of YFP photobleaching and ensure that relatively little mCer3 was inadvertently photobleached.

### Protein Work

Total protein was extracted from 14-day-old plants by grinding in lysis buffer (20 mM Tris, pH 8.0, 150 mM NaCl, 1 mM EDTA, pH 8.0, 1% Triton X-100, 1% SDS with fresh 5 mM DTT, and 100 μM MG132). NF-YA-CFP/HA was probed with high affinity anti-HA primary antibody (cat#11 867 423 001; Roche) and goat anti-rat secondary antibody (cat#SC-2032; Santa Cruz Biotechnology). NF-YC3 and NF-YC4 were detected by previously described native antibodies (51). The Bio-Rad ChemiDoc XRS imaging system was used for visualizing the protein blot after incubations with ECL plus reagent (cat#RPN2132; GE Healthcare). Equivalent loading and transfer efficiency was determined by staining the protein blot with Ponceau S (cat#P3504; Sigma-Aldrich).

### RNA Sequencing and Analysis

Seedlings grown for seven days on B5 media in continuous white light. Total RNA was isolated using the E.Z.N.A. Plant RNA Kit from (Omega Biotek, Cat#R6827). To ensure low levels of contaminating ribosomal RNA, two rounds of poly-A mRNA purification were performed using the μMACS mRNA Isolation Kit (Miltenyi Biotech, Cat#130-090-276). Indexed RNASeq libraries were prepared from 100 ng of poly-A RNA starting material using the NEXTflex Illumina qRNA-Seq Library Prep Kit (Bioo Scientific, Cat#5130). Sequencing of 150 bp paired end reads was performed on an Illumina HiSeq 2500 in rapid output mode at the Texas A&M Agrilife Research Facility (College Station, TX). Sample de-multiplexing was performed using CASAVA software v1.8.2 and bcl2fastq was performed using conversion software v1.8.4.

Resulting sequences were trimmed and quality checked using the pipeline detailed at the iPlant Collaborative Discovery Environment (http://www.iplantcollaborative.org). Sequences were mapped to the TAIR 10 representative gene models set using Burrows-Wheeler Aligner (111, 112) within iPlant. Differential gene expression was determined using the Bioconductor package edgeR (113). Gene Ontology overrepresentation analyses were performed in AmiGO 2 version 2.3.2 (114, 115).

### Image Processing and Figure Construction

All image processing and figure construction was performed in either FIJI, Photoshop (www.adobe.com), or Prism (www.graphpad.com).

## Acknowledgements

We thank Dr. Ben Smith (University of Oklahoma) for assistance with FLIM-FRET measurements and Dr. Min Ni (University of Minnesota) for critical reading of the manuscript. The *cop1-4* mutant allele and *cop1-4 co-9* cross were kindly provided by George Coupland (Max Planck Institute).

## Author Contributions

RWK, ZAM, and BH conceived and managed the project; DP and KKG performed initial analyses of *nf-y* mutants; CLS performed the NF-YA related experiments; JRR and RWK prepared RNASeq samples and libraries; ZAM and RWK performed and analyzed all other experiments; ZAM, RWK, and BFH wrote the manuscript. All authors read and edited the manuscript.

## Supplemental Figure Legends

**Figure S1.**
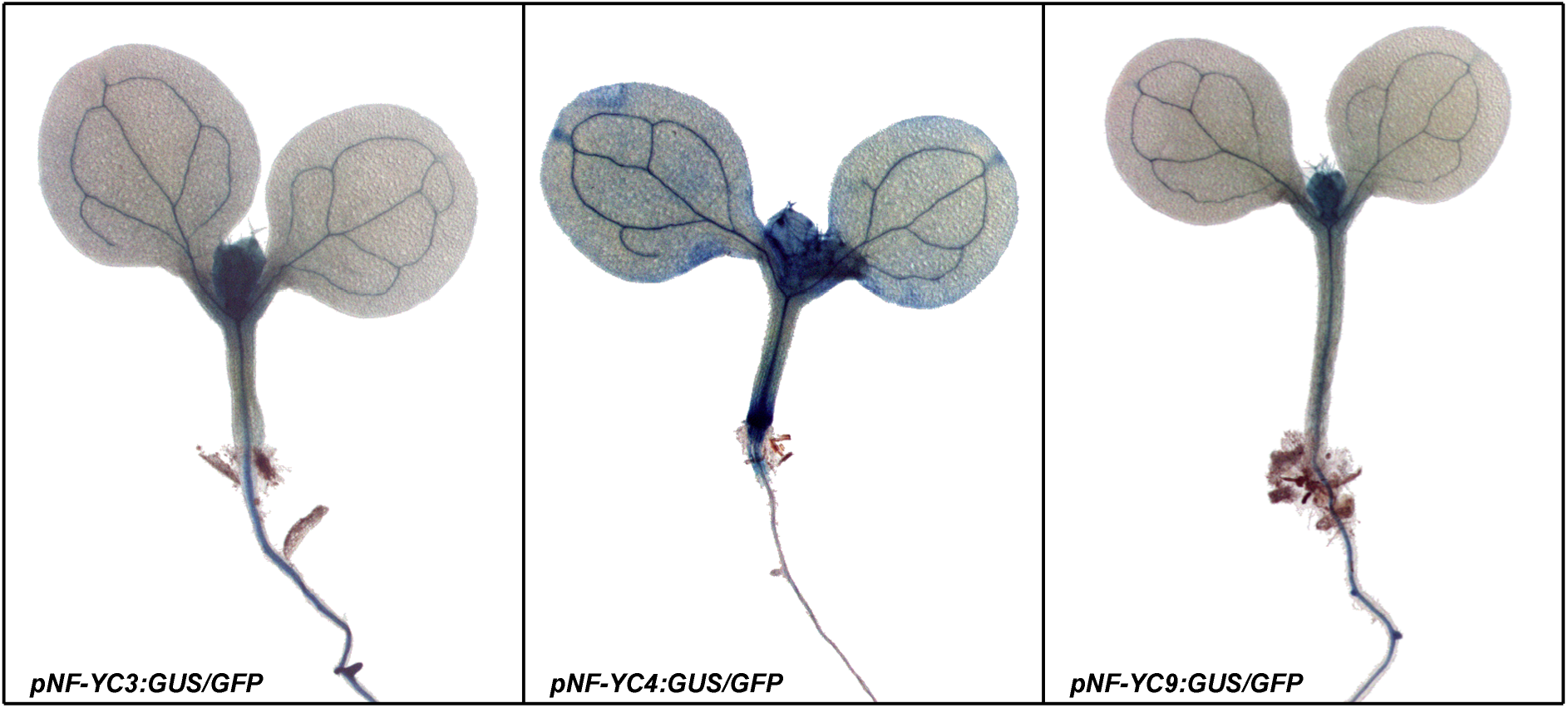
NF-YC3, 4, and 9 are expressed during early seedling development. Promoter-GUS fusions for NF-YC3 (left), NF-YC4 (middle), and NF-YC9 (right) were used to analyze expression patterns in 5-day old plants.

**Figure S2.**
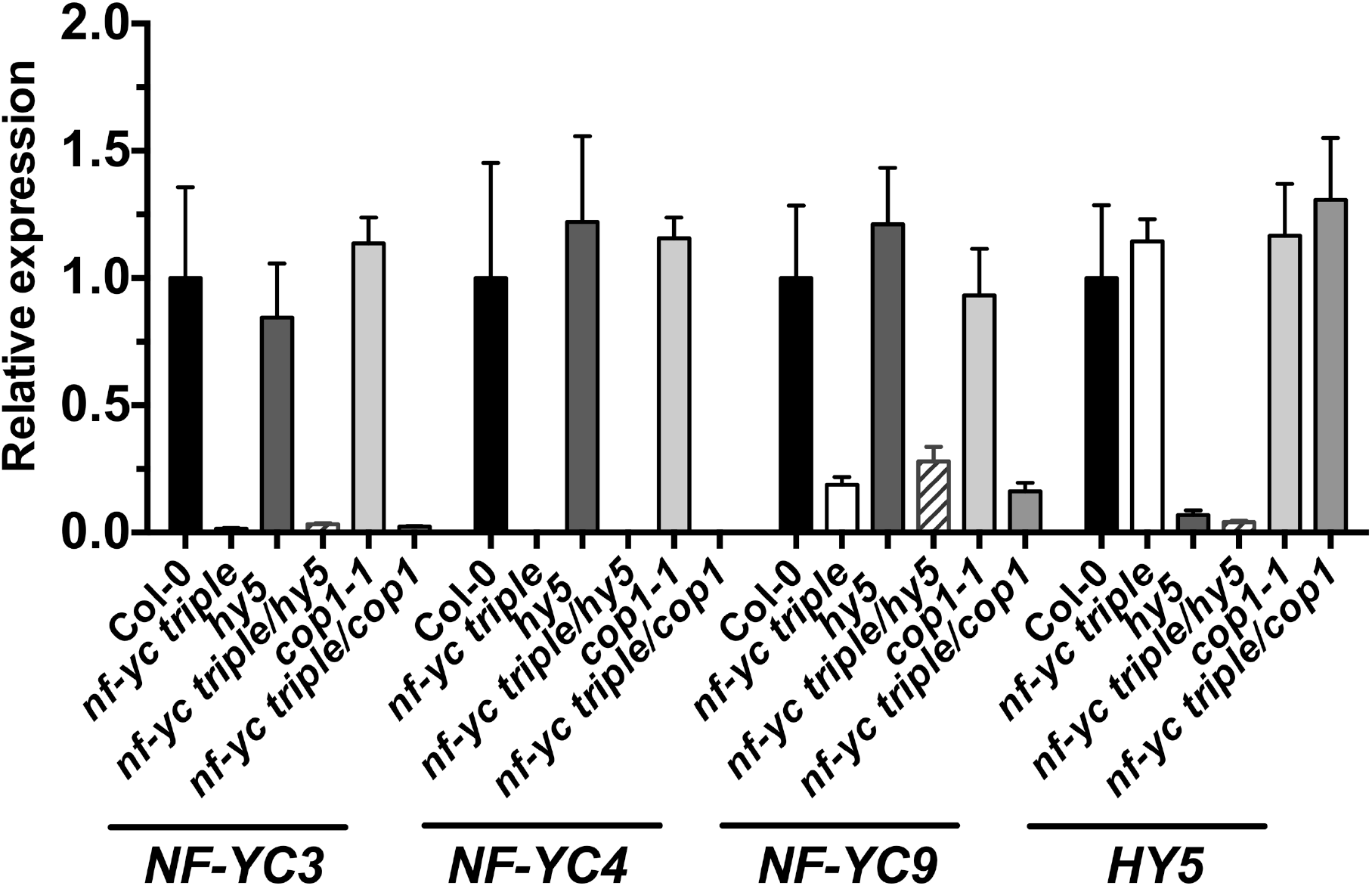
Validation of knock-out and knock-down alleles in higher-order mutants. Expression levels of NF-YC3, NF-YC4, NF-YC9, and HY5 were observed in key genotypes used in this study. Note that NF-YC9 is weakly expressed from the *nf-yc9-1* allele, as previously reported (51). Error bars represent SEM.

**Figure S3.**
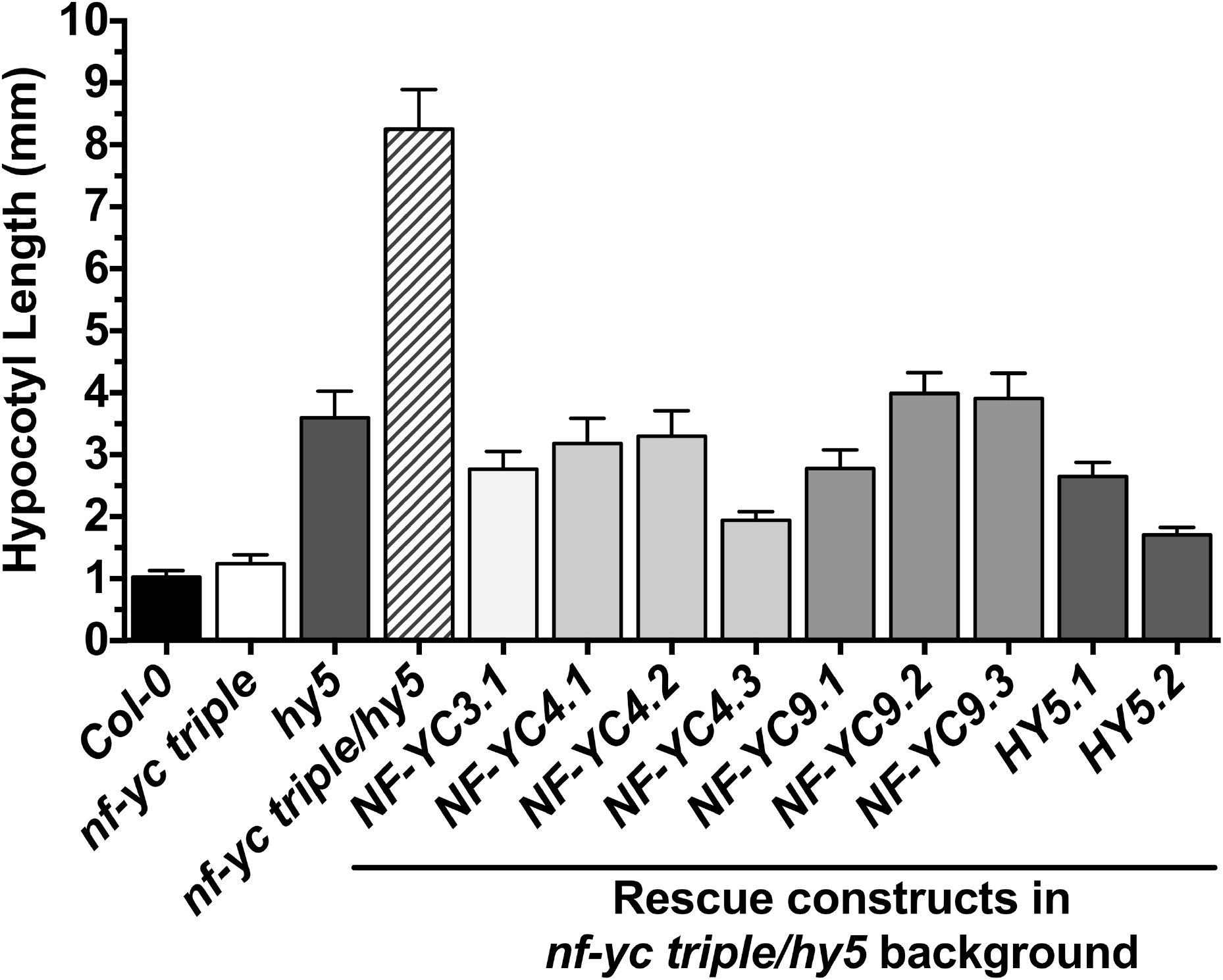
Complementation of the *nf-yc triple hy5* quadruple mutant long hypocotyl phenotype. Native-promoter driven NF-YC3, NF-YC4, and NF-YC9, as well as 35S promoter driven HY5, are able to complement the *nf-yc triple hy5* mutant hypocotyl phenotype in 5 day old, cWL-grown plants. Error bars represent 95% confidence intervals.

**Figure S4.**
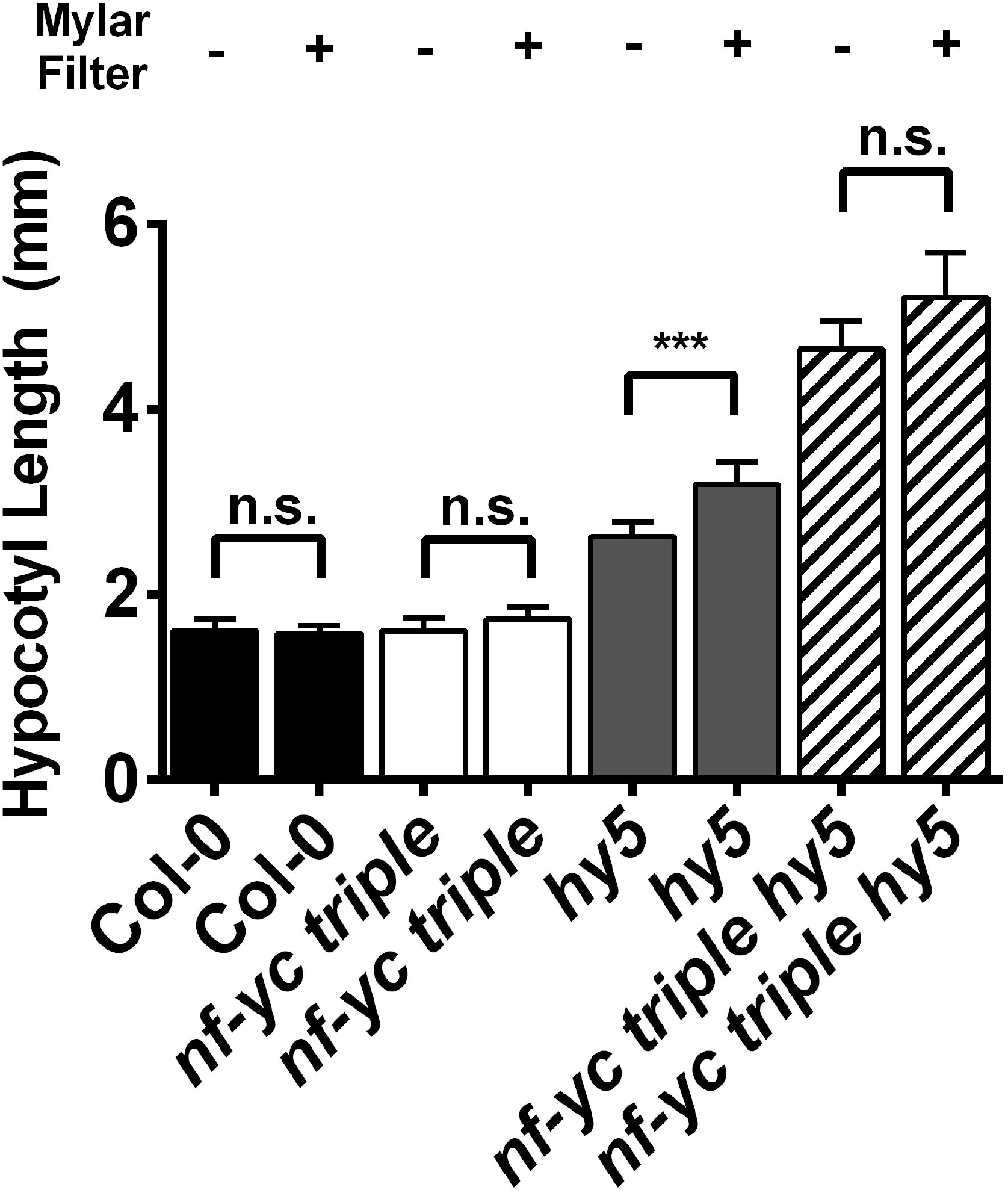
The *nf-yc triple* mutant does not have a hypocotyl elongation response to filtering out UV light. Hypocotyl elongation was assayed in 5 day old, cWL-grown seedlings in the presence or absence of a mylar filter. Error bars represent 95% confidence intervals. Significant differences (and lack thereof) were detected through unpaired t tests (Col-0, *nf-yc triple, hy5*) and an unpaired t test with Welch’s correction for unequal variances (*nf-yc triple hy5*). ***, p < 0.01; n.s., not significantly different.

**Figure S5.**
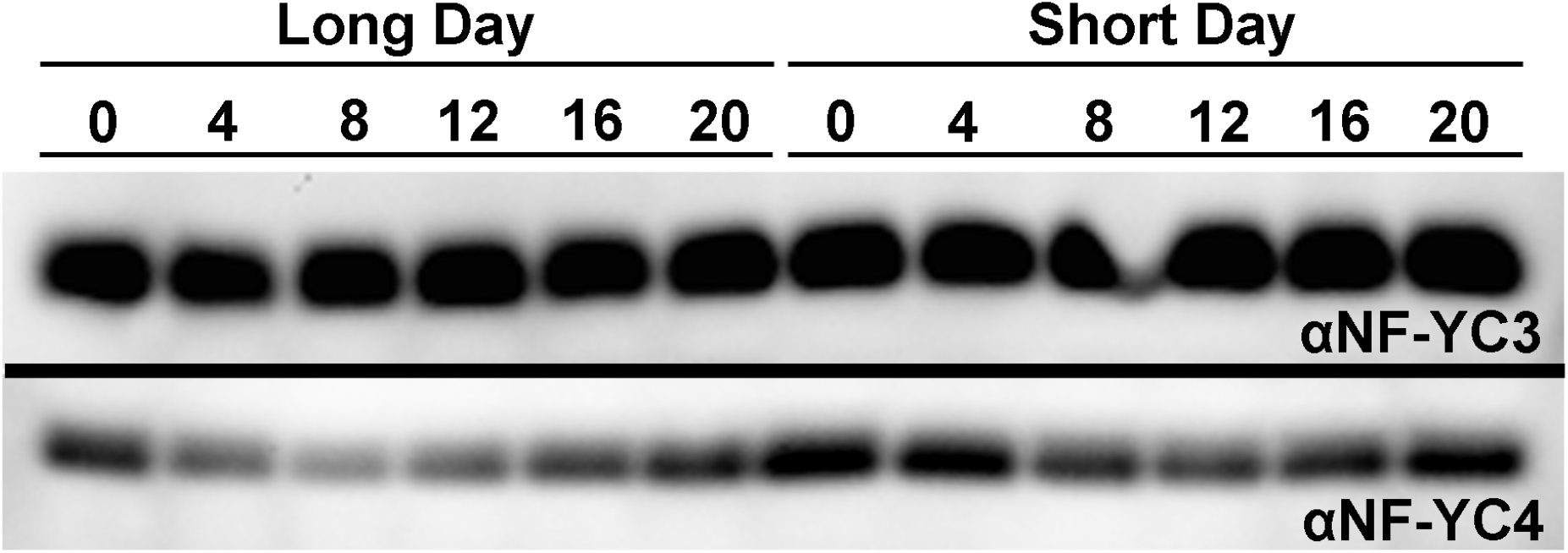
NF-YC3 and NF-YC4 accumulate at relatively steady levels over a 24 hour cycle. Proteins samples were extracted every four hours from both long day (16hr light, 8hr dark) and short day (8hr light, 16hr dark) grown plants. Proteins were detected with previously described, native antibodies (51).

## Supplemental Tables

**Table S1. Lists of all differentially expressed genes identified through RNASeq in *nf-yc triple, hy5*, and *nf-yc triple hy5* mutants**.

**Table S2. GO Enrichment of down-regulated gene sets and regulatory groupings identified through RNASeq**.

**Table S3. GO Enrichment of up-regulated gene sets and regulatory groupings identified through RNASeq**.

**Table S4. Allele designations, descriptions and references for all mutant lines used in this study**.

